# Towards the Development of Isoenergetic Peptide Nucleic Acid Based Probes Targeting Double-Stranded RNAs Through Enhancing Sequence-Specific Stacking Interactions

**DOI:** 10.1101/2025.06.28.662161

**Authors:** Rongguang Lu, Kun Xi, Ruiyu Lin, Manchugondanahalli S. Krishna, Yanyu Chen, Xuan Zhan, Honghao Zeng, Xue Li, Shijian Fan, Hanting Zhou, Wenzheng Wu, Yun Lian, Yuze Dai, Lizhe Zhu, Guobao Li, Gang Chen

## Abstract

Peptide nucleic acid (PNA), a synthetic nucleic acid analog, exhibits substantial potential in biotechnology and therapeutic applications due to its high binding affinity and nuclease/protease resistance. Chemically modified PNAs capable of forming stable triplex structures with double-stranded RNAs (dsRNAs) under near-physiological conditions further expand their utility by enabling sequence-specific precise targeting and probing of functional RNA structural motifs. However, the presence of inverted Watson-Crick pairs (C-G and U-A) may significantly weaken the triplex formation of the dsRNA-binding PNAs (dbPNAs). Our previous work demonstrated that dbPNA P3 (composed of L, T, and Q monomers for the recognition of G-C, A-U, and C-G base pairs, respectively) can stimulate ribosomal frameshifting by binding to rHP2, a model RNA hairpin structure in an mRNA, albeit with suboptimal efficiency, due to its significantly weakened Q•C-G triple formation. We hypothesize that incorporating s^2^U adjacent to Q may offer unique stacking and hydrogen bonding interactions facilitating the development of isoenergetic dbPNA-based probes binding toward dsRNAs with varied sequences. In this study, we investigate how incorporating s^2^U adjacent to Q residues in P3 influences its binding to rHP2. Bio-layer interferometry (BLI) and non-denaturing polyacrylamide gel electrophoresis (PAGE) analyses demonstrate that substituting T with s^2^U at the N-terminal position adjacent to Q (P3-2QT) enhances binding affinity by ∼10-fold compared to unmodified P3, whereas C-terminal substitution (P3-TQ2) yields only a 2-fold improvement. Consistent with these findings, a cell-free dual-luciferase reporter assay reveals that P3-2QT significantly increases ribosomal frameshifting efficiency compared to P3 and P3-TQ2. Molecular dynamics simulations further indicate that P3-2QT maintains enhanced PNA-PNA stacking stability, particularly between s^2^U3 and Q4, suggesting a structural basis for its superior activity. Intriguingly, analogous s^2^U substitution in the P5 oligomer with the Q replaced by L does not confer a comparable enhancement in binding to the target RNA (rHP1) and frameshifting stimulation, highlighting the context-dependent nature of this modification. To assess the broader applicability of the s^2^U-Q motif, we examined its effect in dbPNAs targeting the RNA panhandle structure of influenza virus A and precursor microRNA-21, respectively. Both PAGE and BLI data confirm that s^2^U incorporation improves binding affinity, reinforcing the generality of this strategy. These findings underscore the potential of sequence-dependent uracil thiolation in optimizing triplex-forming dsRNA-binding PNAs, warranting further exploration of modified nucleobase designs to enhance their binding and functional properties.

**Graphical abstract:** 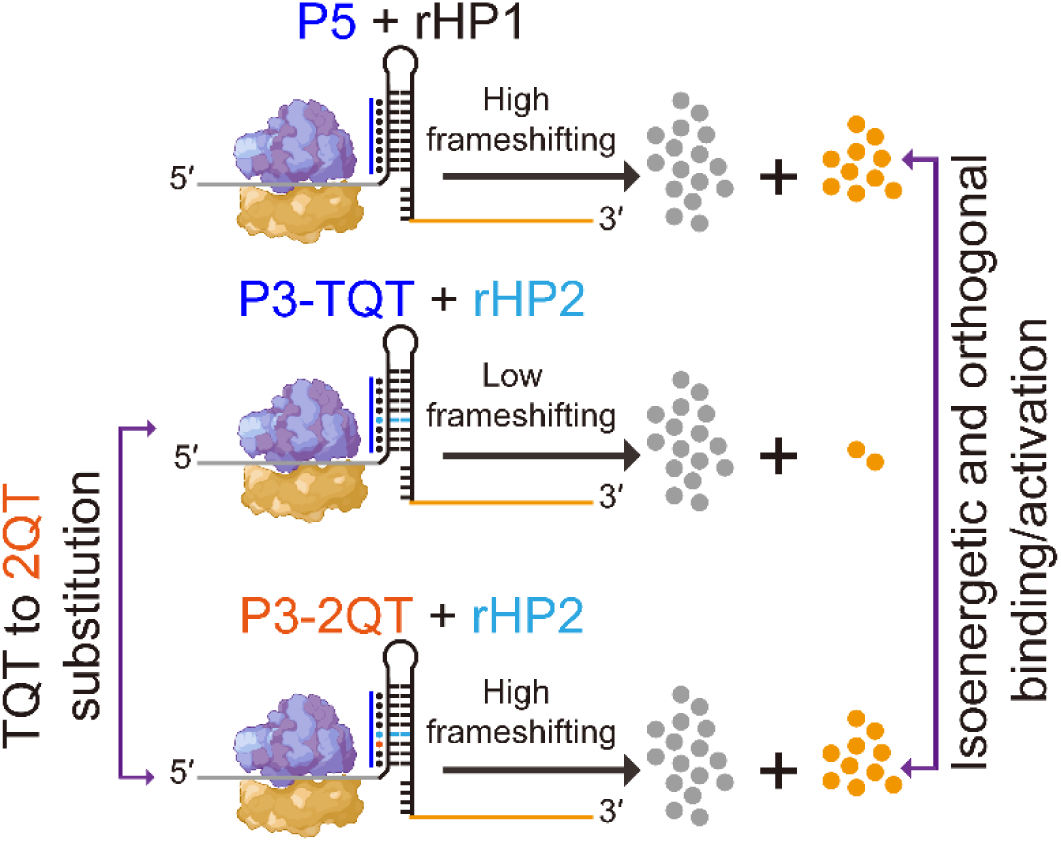

**Highlights:**

1. Thiolation of uracil upstream of Q base in PNA markedly enhances triplex binding affinity and enables isoenergetic and orthogonal targeting/activation.
2. N-terminal s^2^U modification adjacent to Q residues increases PNA-dsRNA binding by ∼10-fold and significantly boosts ribosomal frameshifting efficiency.
3. Biophysical assays and molecular dynamics simulations reveal that thiolation improves PNA stacking stability and slows dissociation, thereby enhancing functional activity toward structured RNA targets.
4. The general applicability of s^2^U modification is demonstrated with influenza A virus RNA panhandle structure and precursor micoRNA-21 targets, underscoring its broad potential for optimizing PNA-based therapeutics and biotechnological tools.

## Introduction

RNA molecules serve as fundamental macromolecules orchestrating diverse biological processes. Their intricate secondary and tertiary architectures govern critical cellular functions, including gene coding, decoding, alternative splicing, and transcriptional/translational regulation ^1,2^. Cutting-edge research elucidating RNA structural thermodynamics, conformational dynamics, and mechanistic functionalities has established a robust framework for pioneering nucleic acid-based innovations ^3,4^. This knowledge has catalyzed the development of numerous RNA-targeted probing and therapeutic modalities, encompassing peptide nucleic acids (PNAs), small interfering RNAs (siRNAs), and small molecule ligands ^5–15^.

Crucially, PNA retains sequence/structure-specific hybridization capabilities, enabling high-affinity binding to single-stranded (ssDNA/ssRNA) via Watson-Crick base pairing or to double-stranded (dsDNA/dsRNA) through Hoogsteen-like interactions, forming stable triplex structures ^9,14,16,17^. These properties position PNA as a versatile programmable platform with applications spanning antisense therapeutics, gene editing, and diagnostic probes ^9–11,14,18–24^. Notably, preclinical studies have demonstrated the biological efficacy of antisense PNAs in animal models, validating their potential for therapeutic intervention ^10,11,13,18,20,22,23^.

Our lab and others have made a series of efforts to create and optimize chemically modified PNA monomers binding to double-stranded RNA (termed dbPNA) through major-groove triple helix formation, including 2-thiouracil (s^2^U), thio-pseudoisocytosine (L) and guanidine-modified 5-methyl cytosine (Q)^9,25–36^ **(Figure 1)**. dbPNAs exhibit high specificity for double-stranded RNA (dsRNA) over single-stranded RNA (ssRNA) or double-stranded DNA (dsDNA). Strategically integrating dbPNA with conventional antisense PNA (asPNA) allows the development of dual-action or dual-affinity PNA (daPNA) for the simultaneous targeting of dsRNA-ssRNA junctions^14^ (cite the CRPS paper). Through programmable design and solid-phase synthesis, these synthetic oligomers demonstrate functional biological activity in both cell-free systems and live cells by targeting structured RNA elements including microRNA precursors, viral RNA motifs, and other functional non-coding RNAs ^14,24,27,37^.

**Figure 1.**
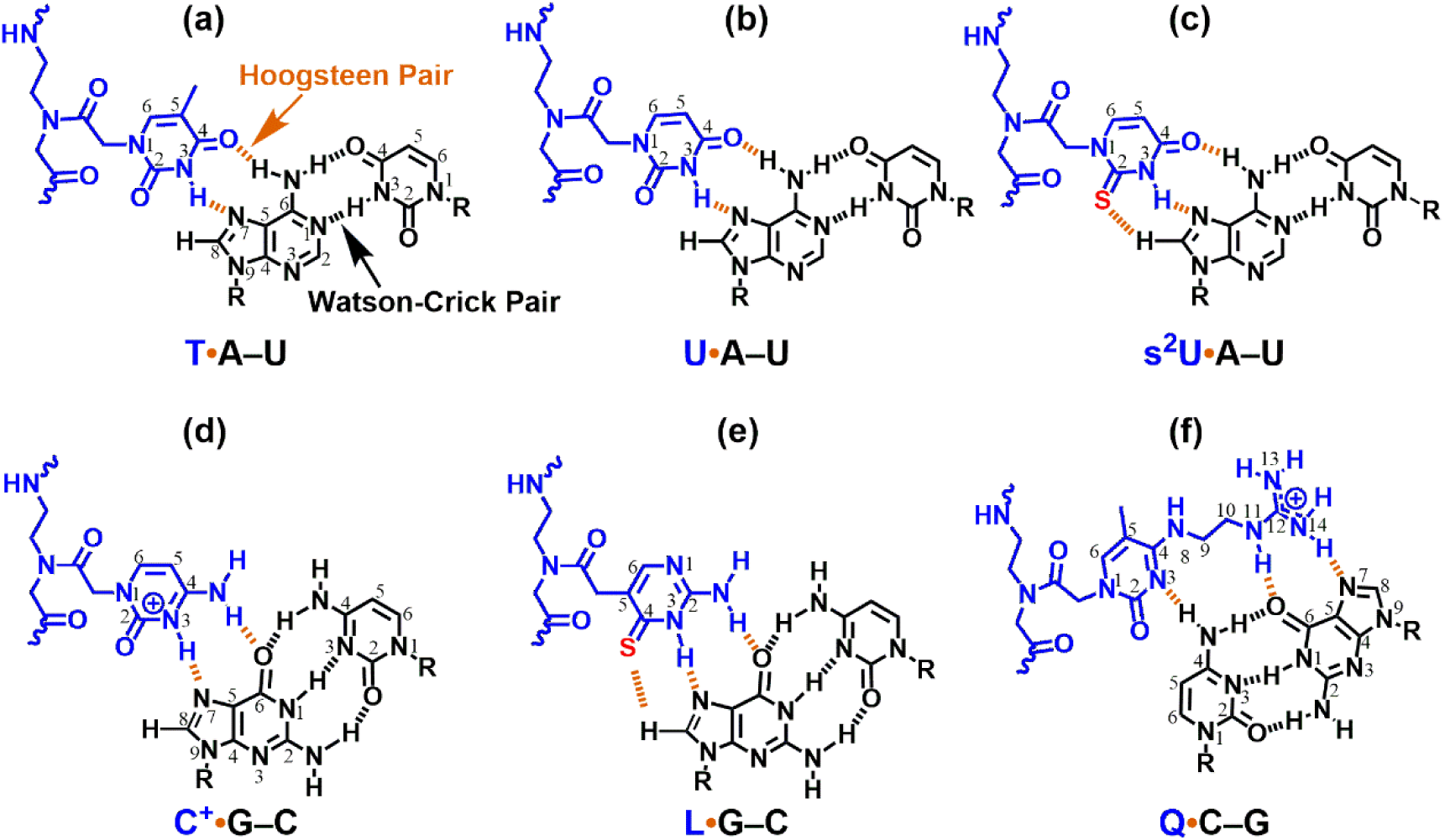
Chemical structures of dbPNA-RNA_2_ triples. Hydrogen bonding patterns for base triples formed between PNA bases (L, Q, s^2^U, U and T): T·A-U (**a**), U·A-U (**b**), s^2^U·A-U (**c**), C^+^·G-C (**d**), L·G-C (**e**), and Q·C-G (**f**) are shown. PNA and RNA are presented in blue and black, respectively. “R” represents the sugar-phosphate backbone of RNA. Orange and black dashed lines indicate hydrogen bonding interactions for PNA with RNA and RNA with RNA, respectively. The sulfur atoms in the s^2^U and L bases are shown in red. In the names of each of the base triples, the Hoogsteen and Watson-Crick pairs are indicated by dots and lines, respectively.

Building on our prior characterization of model RNA hairpins (rHP1 and rHP2, **Figure 2**) for PNA binding and frameshifting modulation ^14,37,38^. This study employs non-denaturing polyacrylamide gel electrophoresis (PAGE) and bio-layer interferometry (BLI) to quantify the binding properties of dbPNA P3 and its s²U-substituted variants with target RNA rHP2 with an inverted C-G pair. The inverted C-G pair can be recognized by a modified PNA base Q through the Q•C-G base triple formation (**Figure 1**), however, often with significantly weakened binding compared to the L•G-C triple. Key parameters (association rate: *k*_on_, dissociation rate: *k*_off_, and equilibrium dissociation constant: *K*_D_) are determined to evaluate the impact of s²U incorporation flanking the Q base on triplex stability. We found that s²U adjacent to Q monomer can enhance the binding of dbPNAs to dsRNAs in a sequence-dependent manner. The work provides the foundation for the development of isoenergetic probes for targeting dsRNAs.

**Figure 2.**
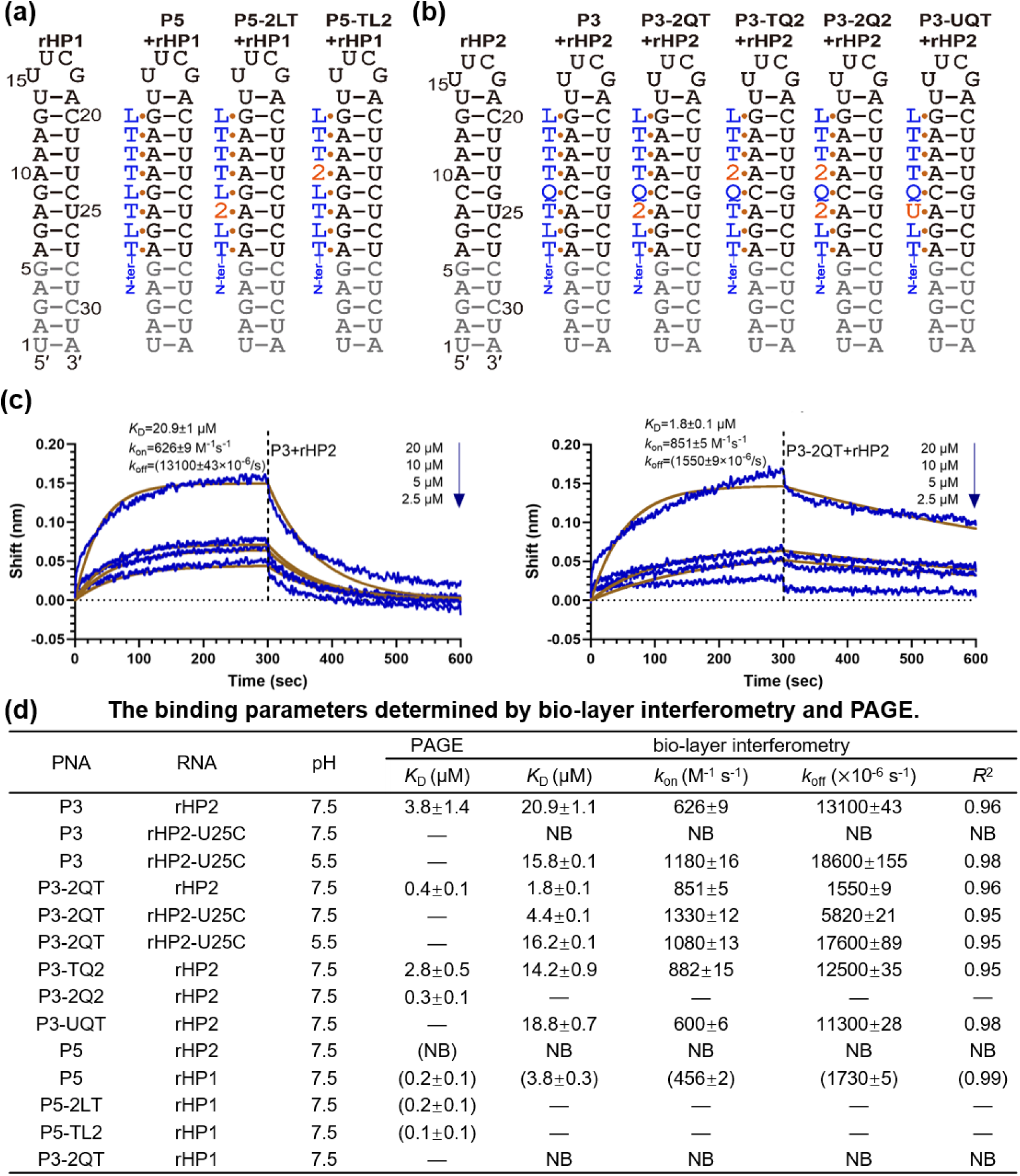
Schematics of PNA-dsRNA triplex formation and binding properties. **(a,b)** Schematics of rHP1 and rHP2 RNAs binding to PNAs. **(c)** Representative binding data of dbPNA P3 and P3-2QT with rHP2 based on bio-layer interferometry assay. **(d)** Binding parameters obtained by nondenaturing PAGE (4℃) and BLI (30℃) in the buffer containing 200mM NaCl, 0.5mM EDTA, 20mM HEPES, pH 7.5. The data shown in the parentheses are from ref ^14,25,27^. NB: no binding observed. “−” indicates that the measurement was not done.

## RESULTS AND DISCUSSION

The PNA monomers Q and L were synthesized following established protocols ^25–27^, while monomers T and s^2^U were commercially procured from QYAOBIO (China). The mass spectrometry results for P3-2QT (2: s^2^U), P3-TQ2, P3-2Q2, P3-UQT, dbPNA-21-TTQT, dbPNA-21-22Q2, IR-2-TQT and IR-2-2Q2 are shown in **Table S1 and Figures S1-S8.** RNA oligonucleotides (sequences cataloged in **Table S2**) were synthesized by Tsingke (China), HPLC-purified, and quantified via UV spectrophotometry. The schematics of PNA•RNA-RNA triples T•A-U, U•A-U, s^2^U•A-U, C^+^•G-C, L•G-C and Q•C-G are shown in **Figure 1**.

Our previous studies demonstrated that P5 (NH₂-lys-TLTLTTTL-CONH₂) and P3 (NH₂-lys-TLTQTTTL-CONH₂) specifically bind to RNA hairpins rHP1 and rHP2, respectively (**Figure 2**). While rHP2 shares structural similarity with rHP1, the key distinction lies in the PNA-targeted base pair at position 4, which changes from G-C (rHP1) to C-G (rHP2) (**Figure 2a,b**). Notably, both s²U-incorporating P5 derivative P5-2LT (NH₂-lys-TL2LTTTL-CONH₂) with s²U at position 3 and P5-TL2 (NH₂-lys-TLTL2TTL-CONH₂) with s²U at position 5) showed negligible or small enhancement of binding affinity **(Figure 2)**^27^. dbPNA P3 (NH_2_-Lys-TLTQTTTL-CONH_2_) can bind to rHP2, but with suboptimal stability^25^. To enhance PNA P3 binding affinity with a goal of developing isoenergetic dsRNA-binding probes, we systematically introduced s^2^U at positions adjacent to Q monomer **(Figure 2b)**: The P3-variant P3-2QT (NH₂-Lys-TL2QTTTL-CONH₂) incorporates s²U at position 3, while P3-TQ2 (NH₂-Lys-TLTQ2TTL-CONH₂) places s²U at position 5. The double-modified P3-2Q2 (NH₂-Lys-TL2Q2TTL-CONH₂) combines both substitutions, and P3-UQT (NH₂-Lys-TLUQTTTL-CONH₂) maintains a standard U substitution at position 3 (**Figure 2b**).

Nondenaturing PAGE and BLI were employed to determine the triplex formation between P3 and rHP2 RNA construct **(Figures 2c, S9)**. The nondenaturing PAGE data revealed a binding affinity (*K*_D_ = 3.8 ± 1.4 μM), consistent with our previously reported value (*K*_D_ = 4.7 ± 0.6 μM) under near-physiological conditions (200 mM NaCl, pH 7.5, **Figures 2d, S9**) ^25^, reaffirming the weak P3-rHP2 interaction. Remarkably, the s²U-modified variant P3-2QT exhibited dramatically enhanced binding (*K*_D_ = 0.4 ± 0.1 μM), representing an 8-fold improvement over unmodified P3 **(Figures 2d, S9)**. In contrast, P3-TQ2 showed more modest (2-fold) enhancement (*K*_D_ = 2.8 ± 0.5 µM). The stabilization effect was also observed in the double-modified P3-2Q2 (*K*_D_ = 0.3 ± 0.1 µM), with s²U substitutions flanking both sides of the Q residue in the sequence **(Figures 2d, S9)**.

Consistent with the PAGE results, BLI analysis further confirmed the enhanced binding affinity of P3-2QT to rHP2, exhibiting a *K*_D_ of 1.8 ± 0.1 μM **(Figure 2c,d)**. This affinity is approximately 10-fold, 7-fold, and 10-fold higher than that of P3, P3-TQ2, and P3-UQT, respectively **(Figures 2c,d, S9, S10)**. Notably, all PNAs demonstrated comparable association rates (*k*_on_): 626 ± 9 M⁻¹s⁻¹ for P3, 851 ± 5 M⁻¹s⁻¹ for P3-2QT, 882 ± 15 M⁻¹s⁻¹ for P3-TQ2, and 600 ± 6 M⁻¹s⁻¹ for P3-UQT when binding to rHP2. In contrast, the dissociation rate constant (*k*_off_) for P3-2QT was markedly reduced at 1,550 ± 9 × 10⁻⁶ s⁻¹, which is 8-fold, 8-fold, and 7-fold slower than those of P3 (13,100 ± 43 × 10⁻⁶ s⁻¹), P3-TQ2 (12,500 ± 35 × 10⁻⁶ s⁻¹), and P3-UQT (11,300 ± 28 × 10⁻⁶ s⁻¹), respectively **(Figures 2c,d, S10)**. These results show that the enhanced binding affinity of P3-2QT is primarily attributable to its reduced dissociation rate.

This observation aligns with the known biophysical properties of s²U: the sulfur atom at the 2-position of uridine promotes more rigid and stable stacking interactions within the nucleic acid duplex or triplex^39,40^. Such stabilization is crucial for maintaining the integrity of higher-order RNA structures under physiological conditions. Moreover, s²U and its C-5 derivatives are predominantly found at the wobble position of tRNAs, where they enhance codon–anticodon pairing fidelity by strengthening hydrogen bonding and base stacking^39,41–43^.

The effect of pH was studied for PNA oligomers P3-2QT, P3-TQ2, and P3-2Q2 at pH 6.0 and 8.0 **(Figure S9)**. The PAGE data show that PNAs incorporated with Q, s^2^U, T and L-monomer do not show significant changes in binding affinity at various pH conditions. P3-2QT binds the target rHP2 with a *K*_D_ 0.4 ± 0.1 µM and 1.0 ± 0.3 µM at pH 6.0 and 8.0, respectively **(Figure S9)**. Whereas P3-TQ2 has a *K*_D_ 2.8 ± 0.5 µM and 7.2 ± 2.8 µM at pH 6.0 and 8.0, respectively, with slightly stronger binding at pH 6.0 over pH 8.0. A similar trend is observed for P3-2Q2 with a binding value of 0.3 ± 0.1 µM, 0.8 ± 0.2 µM at pH 6.0 and 8.0 **(Figure S9)**. The effect of salt concentration was studied using 2 mM MgCl_2_ at pH 7.5. The PAGE experiments show that the PNAs P3-2QT, P3-TQ2, and P3-2Q2 are essentially insensitive to the presence of the salt condition **(Figures S9)**.

Next, we assessed the ribosomal frameshifting efficiency of P3 and its s²U/U-substituted oligomers bound to rHP2^14,37^. **Figure 3a** illustrates the programable −1 ribosomal frameshifting mechanism used by viruses to regulate translation product stoichiometry. The frameshifting stimulatory element (FSE) includes a slippery site (XXX YYY Z), a spacer, and a downstream RNA structure (e.g., pseudoknot or stem-loop). During translation, tRNAs initially decode XXY and YYZ in the 0 frame can slip backward to XXX and YYY in the −1 frame, driven by the downstream RNA’s thermodynamic stability ^38,44,45^.

**Figure 3.**
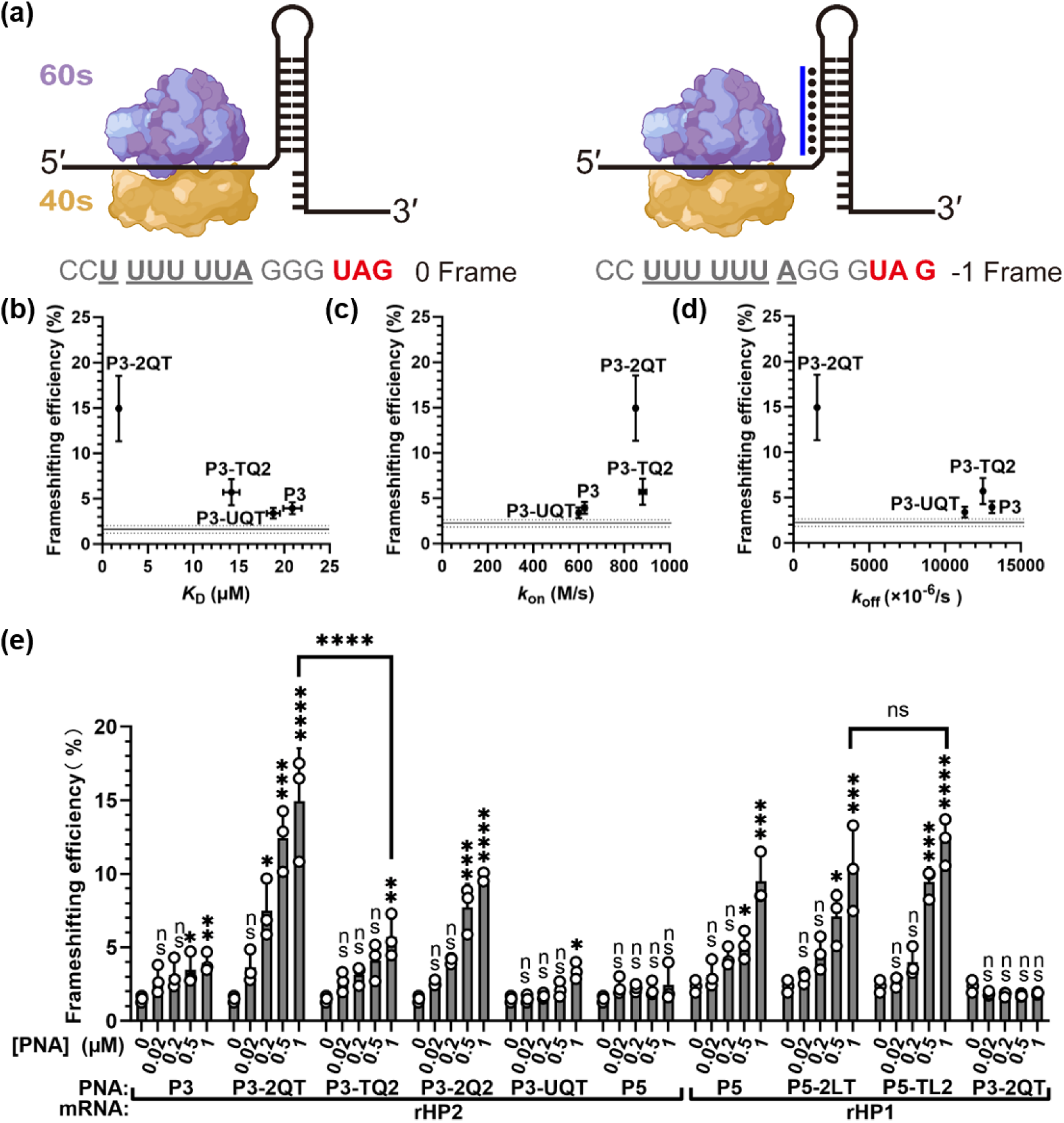
Correlation between binding parameters (*K*_D_, *k*_on,_ and *k*_off_) and frameshifting efficiencies. **(a)** Schematic of ribosomal frameshifting. The binding of a dbPNA (shown in blue) results in an enhanced frameshifting. The slippery site sequences are underlined and the 0-frame stop codon is shown in red. **(b-d)** The *K*_D_ and *k*_off_ values show a positive and inverse correlation with frameshifting efficiency (with the PNA at 1 μM), respectively. Errors of frameshifting efficiencies are reported as standard deviations. The horizontal solid and dashed lines show the average frameshifting efficiencies and standard deviations, respectively. Errors of *K*_D_, *k*_on_ and *k*_off_ values are reported as standard errors. **(e)** Frameshifting efficiency stimulated by PNA-rHP2 triplex formation. The concentration of mRNA is 0.02 μM. The data were analyzed by GraphPad Prism 9.3 and calculated by an ordinary one-way analysis of variance (ANOVA) using Dunnett’s multiple comparisons test against the mean of mRNA alone group. The error bars represent ± S.D. * P 0.05, ** P 0.01, *** P 0.001, **** P 0.0001, ns: not significant. The schematics of the ribosome in panel (a) were drawn by BioRender with permission.

These PNAs stimulate the frameshifting efficiency of rHP2 in a manner that is dependent on their binding affinity **(Figure 3b–d)**. Specifically, the *k*_off_ values are strongly correlated with frameshifting efficiency upon addition of 1 μM PNAs. The frameshifting efficiency of rHP2 increased from 1.5 ± 0.2% (no PNA) to 3.9 ± 0.7%, 5.7 ± 1.4%, and 3.4 ± 0.6% upon addition of 1 μM P3, P3-TQ2, and P3-UQT, respectively **(Figure 3e)**. In contrast, substantially higher frameshifting efficiencies of 14.9 ± 3.6% and 9.7 ± 0.3% were observed with 1 μM P3-2QT and P3-2Q2, respectively **(Figure 3e)**. Consistent with these results, normalized firefly luciferase (Fluc) luminescence increased by 4-fold and 3-fold in the presence of 1 μM P3-2QT and P3-2Q2 compared to the control group with no PNA applied, respectively **(Figure S11)**. In comparison, only a 2-fold increase was observed in the P3 and P3-TQ2 groups **(Figure S11)**. Furthermore, P3-2QT had no significant effect on the frameshifting efficiency or Fluc expression of rHP1 **(Figures 3e, S11)**. Notably, P5-2LT and P5-TL2 did not exhibit significant differences in stimulating frameshifting efficiency or Fluc expression **(Figures 3e, S11)**, suggesting that the pronounced enhancement in activity is specific to PNA oligomers containing a Q monomer with unique stacking interactions involving s²U and Q. Additionally, consistent with our previous findings, P5 neither stimulated frameshifting efficiency nor affected Fluc expression in the context of rHP2 **(Figures 3d, S11)**.

Interestingly, a single-base substitution within the PNA oligomer enables an orthogonal binding between rHP1 and rHP2. Specifically, P5 exhibits strong binding to rHP1, which markedly enhances ribosomal frameshifting and robustly activates downstream −1 frame protein expression. In contrast, P3-TQT binds rHP2 but only induces relatively low frameshifting efficiency. Remarkably, substituting the TQT motif with 2QT in P3 (yielding P3-2QT) results in a 10-fold increase in binding with rHP2 and significantly increases frameshifting efficiency, thereby achieving high levels of activation. Notably, neither P5 with rHP2 nor P3 with rHP1 shows detectable binding or stimulation of frameshifting. This sequence-specific substitution strategy thus enables highly selective, isoenergetic and orthogonal binding and activation between PNA oligomers and their target RNAs, ensuring both the specificity and efficacy^46^. The above experimental binding and frameshifting activity data demonstrate that the binding affinity between PNA and target dsRNA can be substantially enhanced by employing the s^2^U-Q motif within the PNA strand, although the precise binding enhancement mechanism remains unclear. To address this, we conducted unbiased all-atom molecular dynamics (MD) simulations to investigate the stability of three distinct PNAs (i.e., P3, P3-2QT, and P3-TQ2, as depicted in **Figure 4**) binding to the same dsRNA rHP2, thereby elucidating differences in detailed stacking interactions accounting for their different binding affinities. By utilizing recently optimized torsion parameters for the PNA backbone^47^ (cite the ref), we were able to model the PNA-dsRNA complexes without imposing any artificial constraints. Therefore, we propose that, with improved force fields, MD simulations provide a reliable approach for exploring the detailed binding mechanisms of PNA systems. **Figures 4** illustrates the stacking interactions of residues 3 to 5 in P3-related oligomers, aligned with the corresponding RNA base pairs in rHP2. Notably, substitution of T with s^2^U at position 3 in P3-2QT markedly enhances π-π stacking stability by promoting a complete overlap of the pyrimidine rings, probably due the absence of clash between the two methyl groups, present in T3 and Q3 (**Figure 4a,b**). In contrast, this favorable stacking conformation is not observed in P5-2LT or P5-TL2 targeting rHP1 **(Figure S10)**. The stacking arrangement of the upper layer, comprising residues 4 and 5, was found to be similar among P3, P3-2QT, and P3-TQ2 **(Figure 4)**. Thus, through MD simulations, we elucidated the detailed binding modes of these three PNA-dsRNA systems and rationalized the experimental findings by highlighting differences in key hydrogen bonding and π-π stacking interactions.

**Figure 4.**
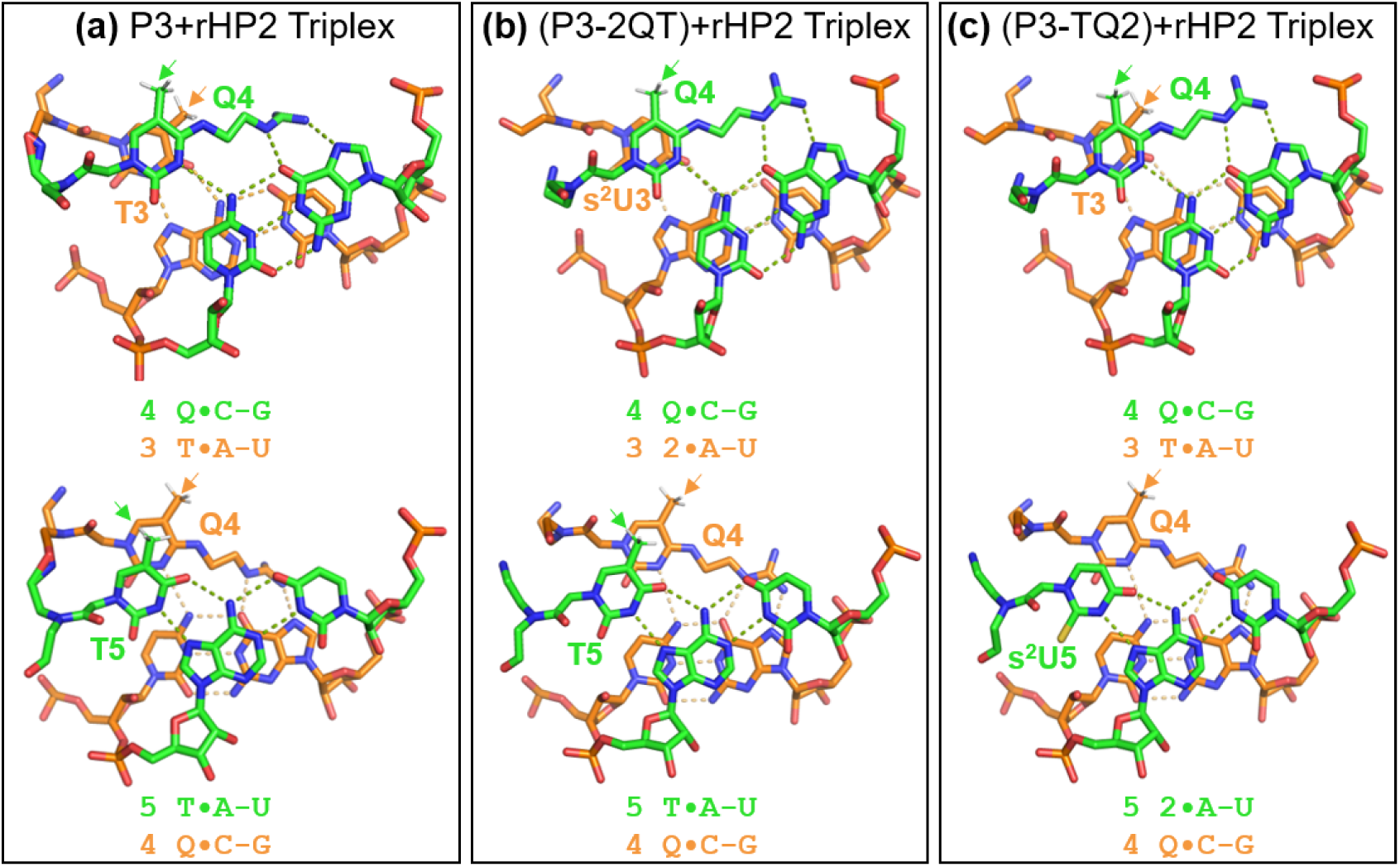
Stacking patterns of PNA·RNA_2_ base triples involving rHP2. The PNA strand is shown on the top left of each panel. (a) Stacking pattern involving residues 3 to 5 of P3 (NH_2_-TLTQTTTL-CONH_2_) targeting rHP2. (b) Stacking pattern involving residues 3 to 5 of P3-2QT (NH_2_-TL2QTTTL-CONH_2_) targeting rHP2. (c) Stacking pattern involving residues 3 to 5 of P3-TQ2 (NH_2_-TLTQ2TTL-CONH_2_) targeting rHP2. The hydrogen atoms of the methyl groups of T and Q are shown and indicated with an arrow. The stacking pattern were visualized by Pymol under orient (Vis) setting.

We subsequently evaluated the generality of this binding mode. We further validated the enhancing effect of s²U substitution on PNA-dsRNA binding using microRNA-21 (miR-21), an oncogenic miRNA that regulates mRNA translation via 3′ UTR binding and requires Dicer-mediated processing of its precursor. Two 7-mer dbPNAs were designed: the unmodified dbPNA-21-TTQT (NH₂-lys-LLQTTQT-CONH₂) and its s²U-substituted variant dbPNA-21-22Q2 (NH₂-lys-LLQ22Q2-CONH₂) **(Figure 5)**. Strikingly, while dbPNA-21-TTQT shows no binding to precursor miR-21 (pre-miR-21), dbPNA-21-22Q2 exhibits strong binding with a *K*_D_ of 623±9 nM **(Figure 5)**. Together with the PH-v example, these results confirm that the s²U-Q modification strategy is broadly applicable for targeting complex functional RNA motifs.

**Figure 5.**
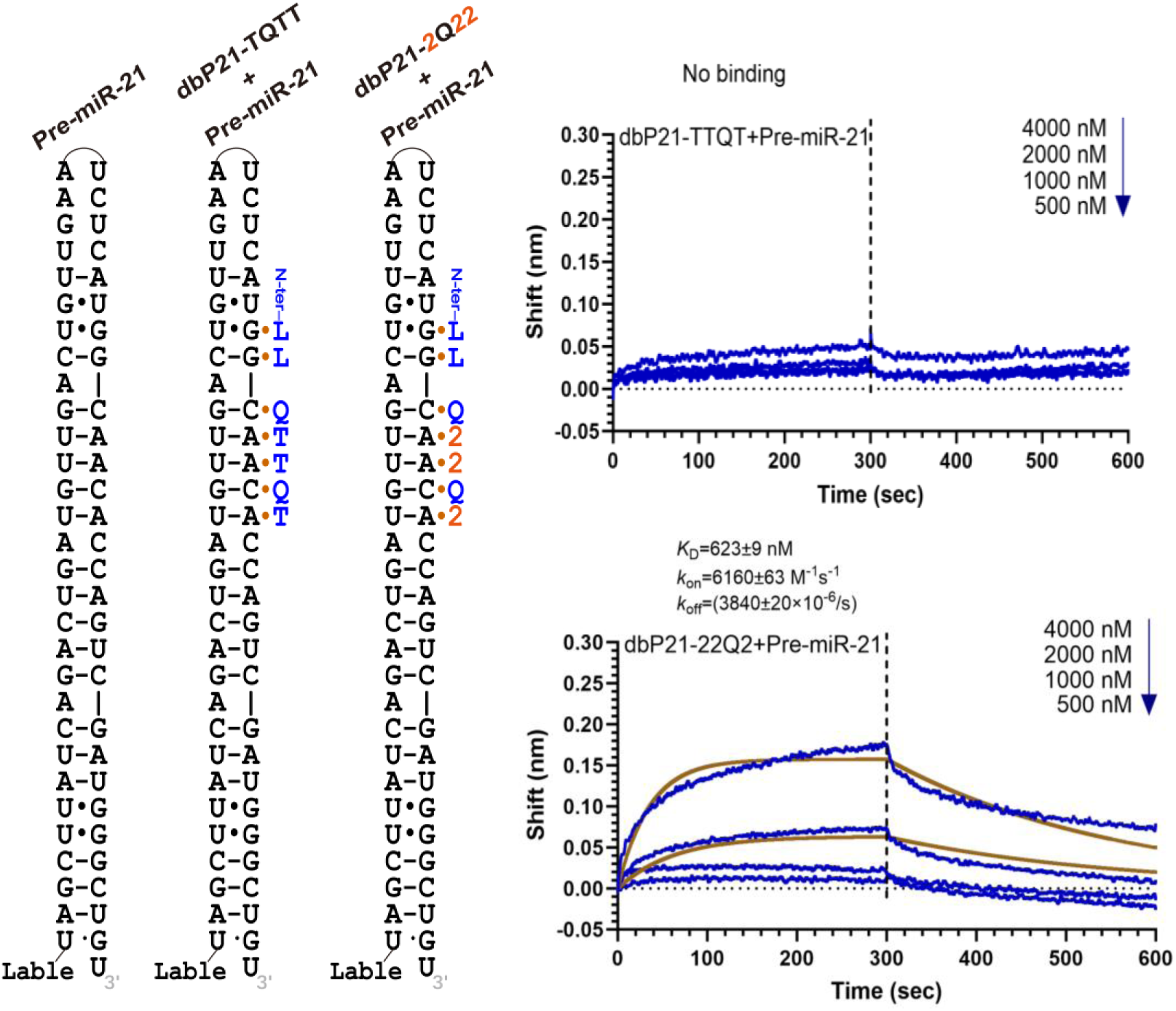
Schematic of dbPNA binding to pre-miR-21 and corresponding BLI data. The left panel shows the secondary structure of pre-miR-21 alone and in complex with dbPNA-21-TTQT or dbPNA-21-22Q2. The right panel presents BLI sensorgrams for the binding of dbPNAs to pre-miR-21. Experimental data are depicted in blue, and the fitting curves are shown in brown. The final concentrations of dbPNA used were 4,000, 2,000, 1,000, and 500 nM from top to bottom.

Previously, we reported that a dbPNA oligomer, IR-1 (NH₂-lys-TLTTTQTLLL-CONH₂), can inhibit influenza A virus (IAV) replication by selectively targeting the viral RNA panhandle structure (PH-v) (24). To assess the applicability of the s^2^U-Q motif in functional RNA structures, we designed IR-2 (NH₂-lys-TLTTTQTLL-CONH₂), a dbPNA variant lacking one C-terminal L monomer compared to IR-1 **(Figure S13a–c)** (24), and substituted both T residues adjacent to the Q monomer with s^2^U, yielding IR-2-2Q2 (NH₂-lys-TLTT2Q2LL-CONH₂). Non-denaturing PAGE data demonstrate that IR-2-2Q2 exhibits significantly enhanced binding to PH-v, with a *K*_D_ of 165 ±76 nM, compared to IR-2-TQT (*K*_D_ = 255 ± 121 nM) **(Figures S13d, S14)**. Consistent with these results, BLI analysis reveals that IR-2-2Q2 (*K*_D_ = 90 ± 1 nM) displays approximately two-fold stronger binding to PH-v than IR-2-TQT (*K*_D_ = 210 ± 1 nM) (**Figure S13e, f**). Notably, IR-2-2Q2 exhibits a two-fold slower dissociation rate (*k*_off_) relative to IR-2-TQT, while the association rates (*k*_on_) remains similar **(Figure S13e, f)**. These findings indicate that substituting T with s^2^U adjacent to the Q monomer in both the P3 and IR-2 series dbPNA oligomers enhances binding affinity to RNA constructs primarily by reducing the dissociation rate.

Due to the presence of an A•C wobble pair in the PH-v structure targeted by IR-2-TQT, which may compromise the affinity enhancement typically conferred by the s^2^U-Q motif, we sought to determine the effects of the A•C wobble pair on triplex formation with P3 and P3-2QT. To this end, a mutant rHP2-U25C was generated. BLI data reveal that P3 does not bind to rHP2-U25C (**Figures 2d, S15**). However, P3-2QT is still able to bind to rHP2-U25C, albeit with a weaker affinity (*K*_D_ = 4.4 ± 0.1 μM) compared to rHP2 (*K*_D_ = 1.8 ± 0.1 μM) under buffer condition (200 mM NaCl, 0.5 mM EDTA, 20 mM HEPES, pH 7.5). Interestingly, the dissociation rate of P3-2QT from rHP2-U25C increased significantly, by approximately threefold compared to rHP2, to 5820 ± 21 × 10⁻⁶ s⁻¹ (**Figures 2d, S15**). This provides an explanation for why IR-2-2Q2 only enhanced binding by two-fold, in contrast to the tenfold increase observed with P3-2QT and the target RNA. Surprisingly, under more acidic conditions (pH 5.5), both P3 and P3-2QT were able to bind to rHP2-U25C with similar *K*_D_ values (**Figures 2d, S15**). It is possible that lowering pH strengthening the wobble A•C wobble pair formation may restrict the flexibility of A for pairing with PNA bases T or s^2^U.

All-atom molecular dynamics simulations were performed to explore the detailed stacking patterns of P3 and P3-2QT with rHP2-U25C, in which the formation of an A•C wobble pair was enforced^38^. The results show that the stacking model of P3 with rHP2-U25C is similar to that with rHP2 (**Figure 4a, Figure S16**). In contrast, P3-2QT exhibits significant differences in stacking with rHP2-U25C compared to rHP2, particularly at the third and forth position, where the stacking of s^2^U and Q is no longer in the complete overlap mode **(Figures 4b, S16)**. This change in stacking geometry may explain the reduced binding affinity observed.

### Conclusions

In this study, we systematically investigated the relationship between PNA stacking interactions— including those involving T, s^2^U, L, and Q monomers—and their binding affinity to RNA constructs, as well as the resulting biological activity in cell-free systems. Our findings demonstrate that uracil thiolation (as in s^2^U) can enhance stacking between the PNA and RNA structure, particularly when positioned at the N-terminal residue adjacent to the Q monomer. Moreover, the incorporation of s^2^U into dbPNA oligomers significantly improves both their binding affinity to RNA constructs and their ability to stimulate ribosomal frameshifting. Collectively, these results establish a novel design principle for isoenergetic dbPNA oligomers, substantially increasing the affinity for structured dsRNA targets with C-G inversions—an advance with important implications for the probing and modulation of RNA function in human diseases.

## MATERIALS AND METHODS

### Solid phase synthesis of PNA oligomers

The unmodified N-(2-aminoethyl) glycine PNA (aegPNA) monomers were procured from ASM Research Chemicals and QYAOBIO (China), while the L- and Q-modified monomers were synthesized following established literature procedures ^25,26^. Oligomer synthesis was conducted on MBHA·HCl-functionalized polystyrene resin with an initial loading of 1.5–1.7 mmol/g, systematically reduced to 0.35 mmol/g through acetic anhydride capping. Solid-phase assembly employed Boc-chemistry with PyBop/DIPEA coupling agents, followed by resin cleavage using a sequential TFA-TFMSA protocol. Crude products underwent ice-cold diethyl ether precipitation, aqueous dissolution, and RP-HPLC purification (water-acetonitrile-0.1% TFA mobile phase). LC-MS/MS characterization confirmed oligomer integrity, with molar extinction coefficients (260 nm) quantified as 8.8 (T), 10.2 (s²U), and 7.3 mM⁻¹cm⁻¹ (Q/L).

### Nondenaturing polyacrylamide gel electrophoresis

Non-denaturing 12% polyacrylamide gel electrophoresis was performed under four distinct buffer conditions: 200mM NaCl, 0.5mM EDTA, 20mM MES, pH 6.0; 200mM NaCl, 0.5mM EDTA, 20 mM HEPES, pH 7.5; 200mM NaCl, 0.5mM EDTA, 2mM MgCl_2_, 20 mM HEPES, pH 7.5; 200mM NaCl, 0.5mM EDTA, 20 mM HEPES, pH 8.0. For Cy3-labeled rHP2-Cy3 analysis, samples containing 0.05 µM RNA were titrated with PNA concentrations ranging from 0 to 10 µM (0, 0.005, 0.01, 0.02, 0.05, 0.1, 0.2, 0.4, 0.7, 1, 1.5, 2, 5, 10 µM) in 20 µL reaction volumes. RNA hairpins were snap-cooled by rapid transfer from 95°C to an ice bath, while PNA-RNA complexes were annealed via slow cooling from 65°C to room temperature. All samples were equilibrated overnight at 4°C prior to electrophoresis. Before loading, samples were supplemented with 35% glycerol (20% of total loading volume) and briefly vortexed. Electrophoresis was conducted in 1× TBE buffer (pH 8.3) at 4°C for 6 hours under constant voltage (250 V).

### Bio-layer interferometry

The binding kinetics of PNAs to biotinylated RNA constructs were analyzed by bio-layer interferometry (BLI) using a Gator® Prime instrument (China) at 30°C, with 300-second association and dissociation phases. Streptavidin (SA) biosensors were employed to immobilize RNA constructs diluted to 0.2 µM in binding buffer (200 mM NaCl, 0.5 mM EDTA, 20 mM HEPES, pH 7.5). PNA samples (excluding P3-TQ2) were serially diluted to 20, 10, 5, and 2.5 µM in the same buffer, while P3-TQ2 concentrations were adjusted to 16, 8, 4 and 2 µM. For assays targeting PH-v RNA with IR-2 and IR-2-2Q2 PNAs, the RNA was annealed by heating to 95°C for 5 min followed by snap-cooling on ice, and PNA concentrations were tested at 5,000, 2,500, 1,250, and 625 nM. Binding parameters (*K*_D_, *k*_on_ and *k*_off_) were derived from both Global and unlinked fitting models. Real-time binding curves (Δλ shift vs. time, 0.1-s resolution) and fitted plots were processed using GraphPad Prism 10 to generate high-resolution figures.

### Plasmid construction

The frameshifting efficiency was quantified using the p2luc dual-luciferase reporter plasmid, generously provided by Prof. Samuel E. Butcher ^45^, wherein the rHP1 and rHP2 DNA sequences were inserted between the Renilla and firefly luciferase coding regions via BamHI and SacI restriction sites, as previously established ^14,37^. A modified in-frame control (IFC) plasmid was generated by disrupting the slippery sequence to enforce 100% translational fidelity^37^. All plasmids were purified using the Monarch Plasmid Miniprep Kit (NEB) followed by ethanol precipitation, with DNA quality and concentrations verified by a NanoDrop One spectrophotometer (Thermo Scientific).

### DNA linearization, transcription and RNA purification

Linearized DNA was generated via polymerase chain reaction (PCR) using primers p2luc-631 and p2luc3431 (Table S2)^14^ and the KOD-Plus-Neo kit (TOYOBO). Each 50 µL reaction contained 5 µL 10× KOD-Plus-Neo Buffer, 5 µL 2 mM dNTPs, 3 µL 25 mM MgSO₄, 1.5 µL each of 10 pmol/µL forward and reverse primers, 2.5 µL 10 ng/µL DNA template, 1 µL 1.0 U/µL KOD-Plus-Neo polymerase, and 30.5 µL nuclease-free H₂O (Sangon). Reactions were mixed thoroughly and cycled in a BioRad thermocycler under the following conditions: 95°C for 30 s; 35 cycles of 95°C for 30 s, 57°C for 30 s, and 72°C for 2 min; followed by a final 72°C elongation for 2 min. PCR products were ethanol-precipitated, quantified via NanoDrop (Thermo Scientific), and transcribed using the HiScribe T7 High-Yield RNA Synthesis Kit (NEB). After 2 h of transcription at 37°C, DNA templates were digested with DNase I (NEB), and RNA transcripts were purified using an RNA cleanup kit (NEB) and resuspended in nuclease-free H₂O for downstream applications.

### *In vitro* translation, dual-luciferase reporter assay and data analysis

The cell-free translation system was adapted from established methods^14,37^ with minor modifications. Briefly, mRNA transcripts (final concentration: 0.02 μM) were pre-mixed with serial dilutions of dbPNA (final concentration: 0, 0.02, 0.2, 0.5, 1 μM) in DEPC-treated water (Sangon). Each 12.5 μL reaction consisted of 8.75 μL nuclease-treated rabbit reticulocyte lysate (RRL, Promega), 1.25 μL PNA solution, and 2.5 μL mRNA, incubated at 30°C for 90 min in a thermal cycler. For luminescence quantification, 2 μL of translation products were combined with 50 μL Dual-Glo Firefly Luciferase Substrate (Promega), and relative luminescence units (RLU) were immediately measured using an EnVision Multimode Plate Reader (PerkinElmer). Subsequently, 50 μL Dual-Glo Stop & Glo *Renilla* Substrate (Promega) was added to quantify RLU for normalization. All conditions were tested in triplicate, with data analyzed in GraphPad Prism. The formula for calculating the frameshifting efficiency is^48^:

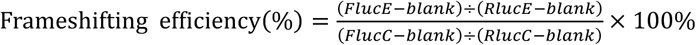

### Structure Modeling

The initial structure of the PNA-dsRNA triplex was built based on the crystal structure (PDB ID: 1PNN) of a PNA·DNA-PNA triplex see the details in Ref ^36,49^. For the Q residue of PNA, its initial structure was generated by Gaussian View, and the initial structure of the s^2^U of PNA was generated by modifying the T residue, i.e., replacing the CH_3_ group with hydrogen and the O_2_ with S_2_. Both initial structures were optimized at the MP2 6-31+G(d)21, 23-26 level using Gaussian 16 (Gaussian, Inc., Wallingford, CT, USA). Then, the PNA strands were capped by adding an acetyl (COCH_3_) group to the N-terminus and an amide (NH_2_) group to the C-terminus. While for the dsRNA, the initial structure was built based on Macromolecular Builder (MMB). After that, the restrained electrostatic potential charges were determined using an antechamber program^50^, and the force field parameters were described using a Generalized Amber Force Field (GAFF) force field. In addition, the recently improved torsion parameters for the backbone of PNA were also employed^47,51^. Then the four PNA-dsRNA MD simulation systems were built. We calculated the root mean square deviation (RMSD) of the central X_3_-Q_4_-X_5_ (X=T or s^2^U) region (highlighted with cyan color) for the PNA-dsRNA complex. As the system quickly reached equilibrium within ∼20 ns, we only used the last 200 ns MD trajectories to analyze the detailed interactions.

## ASSOCIATED CONTENT

### Supporting information

Supplementary data to this article can be found online.

## Author Contributions

Rongguang Lu, Kun Xi, Ruiyu Lin, Manchugondanahalli S. Krishna Yanyu Chen and Xuan Zhan contributed equally to the work.

## Notes

There are no conflicts of interest to declare.

## ACKNOWLEDGMENTS

This work was supported by National Natural Science Foundation of China (NSFC) project (Grant 22177098 to G.C.), The Chinese University of Hong Kong, Shenzhen (CUHK-Shenzhen) University Development Fund (to G.C.), fund from Shenzhen-Hong Kong Cooperation Zone for Technology and Innovation (HZQB-KCZYB-2020056 to G.C.), Shenzhen Third People’s Hospital Research Fund (24250G1021) and China Postdoctoral Science Foundation funded project (2023M742416), Shenzhen Clinical Research Center for Tuberculosis (20210617141509001), Shenzhen Science, Technology and Innovation Committee for the Shenzhen Key Laboratory Scheme (ZDSYS20220507161600001), Guangdong Provincial Basic and Applied Basic Research Fund Project-Youth Funding (2022A1515110577 to X.Z.). This work was also supported by the Warshel Institute for Computational Biology Funding from Shenzhen City and Longgang District, the National Science Foundation of China [Grant 31971179; 32471296], the Science, Technology and Innovation Commission of Shenzhen Municipality [Grant JCYJ20200109150003938, RCYX20200714114645019] and National Supercomputer Center in Guangzhou. We are also grateful to Qiushi Guo for her invaluable support in mass spectrometry data acquisition and interpretation.

## Notes

### Competing Interest Statement

The authors have declared no competing interest.

