## Supporting information for "Towards the Development of Isoenergetic Peptide Nucleic Acid Based Probes Targeting Double-Stranded RNAs Through Enhancing Sequence-Specific Stacking Interactions"

#### Table of Contents


1. **Table S1 and Figure S1 to S8:** Mass spectrum results of PNAs. **S3-S7**

2. **Table S2:** RNA and DNA oligonucleotides used in this study. **S8**

3. **Figure S9:** PAGE characterization of PNAs designed for targeting rHP2. **S9**

4. **Figure S10:** BLI data for PNAs binding to rHP1 and rHP2. **S10**

5. **Figure S11:** Normalized firefly and *Renilla* luciferase RLU of mRNAs rHP1 and rHP2. **S11**

6. **Figure S12:** Stacking patterns of PNA·$RNA_2$ base triples involving rHP1 and PNA segment TLT/2LT/TL2. **S12**

7. **Figure S13:** Schematic of dbPNA binding to PH-v and corresponding PAGE and BLI data **S13**

8. **Figure S14:** Fitting curve of PAGE results for dbPNAs binding to PH-v **S13**

9. **Figure S15:** BLI data for PNAs targeting rHP2-U25C **S14**

10. **Figure S16:** Stacking patterns of PNA·$RNA_2$ base triples for PNAs targeting to rHP2-U25C **S15**

11. **References** **S15**

**Table S1**: Mass spectrum data of PNA sequences.

| PNA | Oligomer Sequence | Chemical Formula | Calculated MW | Obtained MW |
|---|---|---|---|---|
| P3 | $H_2N$-Lys-TLTQTTTL-$CONH_2$ | $C_{95}H_{133}N_{41}O_{28}S_2$ | 2361.46 | (2360.93) |
| P3-2QT | $H_2N$-Lys-TL2QTTTL-$CONH_2$ | $C_{94}H_{131}N_{41}O_{27}S_3$ | 2363.52 | 2361.94 |
| P3-TQ2 | $H_2N$-Lys-TLTQ2TTL-$CONH_2$ | $C_{94}H_{131}N_{41}O_{27}S_3$ | 2363.52 | 2361.94 |
| P3-2Q2 | $H_2N$-Lys-TL2Q2TTL-$CONH_2$ | $C_{93}H_{129}N_{41}O_{26}S_4$ | 2365.56 | 2363.89 |
| P3-UQT | $H_2N$-Lys-TLUQTTTL-$CONH_2$ | $C_{94}H_{131}N_{41}O_{28}S_2$ | 2345.95 | 2345.96 |
| daPNA-21-TTQT | $H_2N$-Lys-LLQTTQT-$CONH_2$ | $C_{87}H_{127}N_{41}O_{23}S_2$ | 2177.95 | 2177.94 |
| daPNA-21-22Q2 | $H_2N$-Lys-LLQ22Q2-$CONH_2$ | $C_{84}H_{121}N_{41}O_{20}S_5$ | 2183.83 | 2183.84 |
| IR-2-TQT | $H_2N$-Lys-TLTTTQTLL-$CONH_2$ | $C_{105}H_{146}N_{46}O_{30}S_3$ | 2627.05 | 2627.05 |
| IR-2-2Q2 | $H_2N$-Lys-TLTT2Q2LL-$CONH_2$ | $C_{103}H_{142}N_{46}O_{28}S_5$ | 2630.97 | 2630.97 |

The oligomer P3 was previously studied and characterized (Toh et al., 2016). “2” represents a 2-thiouracil monomer in the sequences.

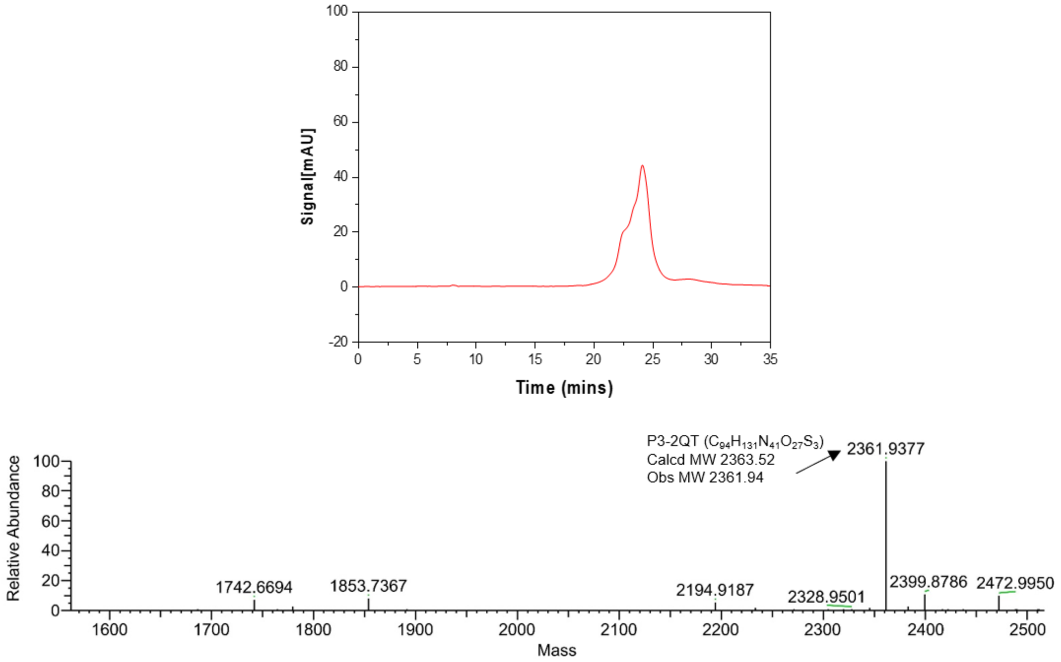


**Figure S1. HPLC and LC/MS data of P3-2QT.**

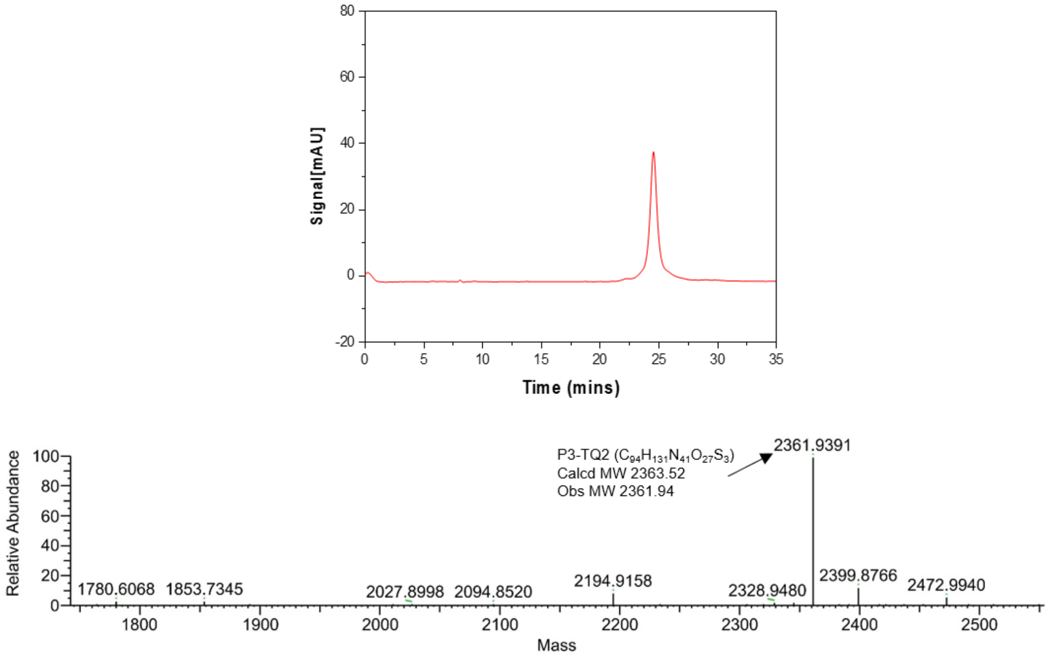


**Figure S2. HPLC and LC/MS data of P3-TQ2.**

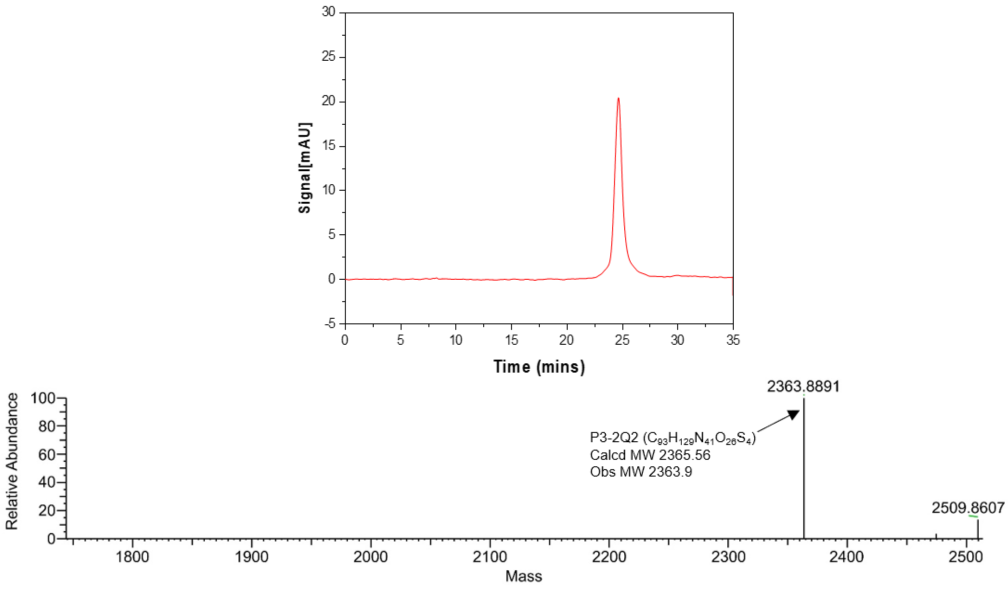


**Figure S3. HPLC and LC/MS data of P3-2Q2.**

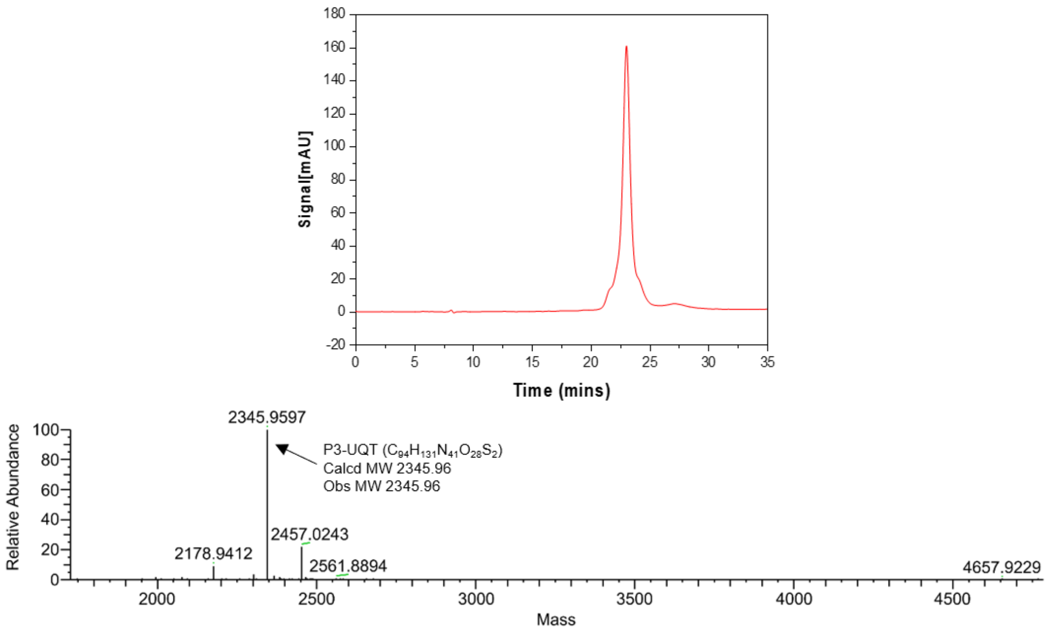


**Figure S4. HPLC and LC/MS data of P3-UQT.**

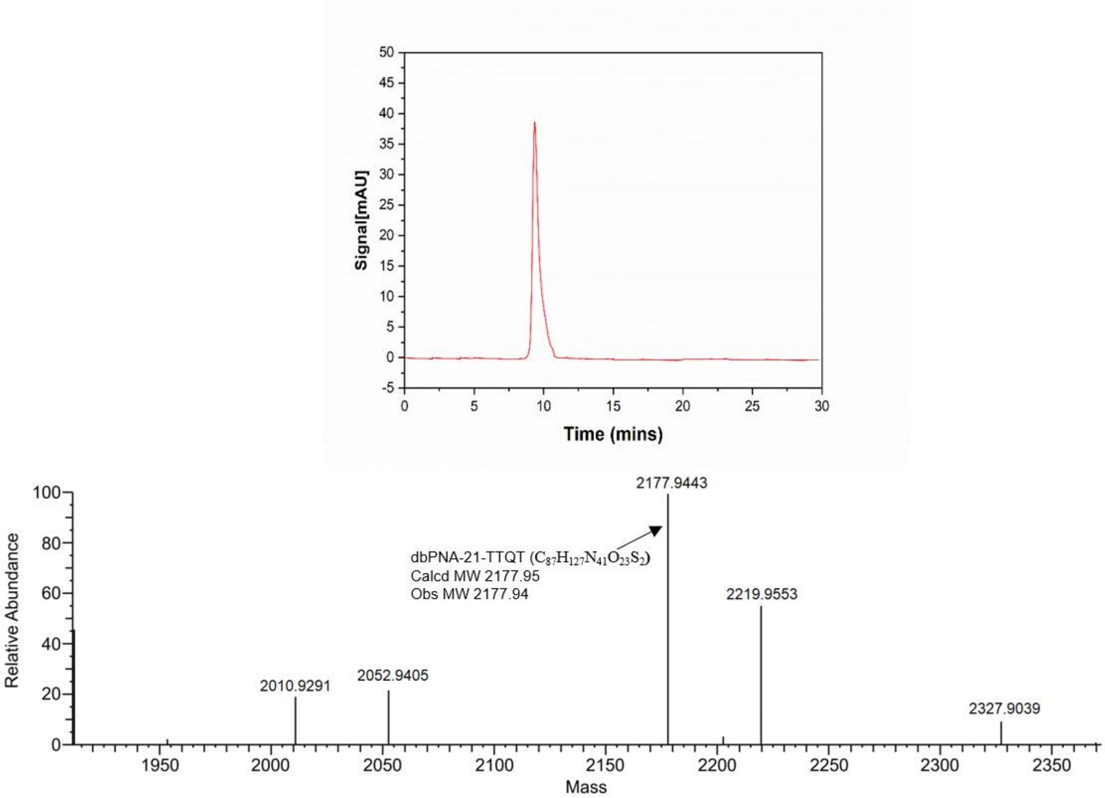


**Figure S5. HPLC and LC/MS data of daPNA21-TTQT.**

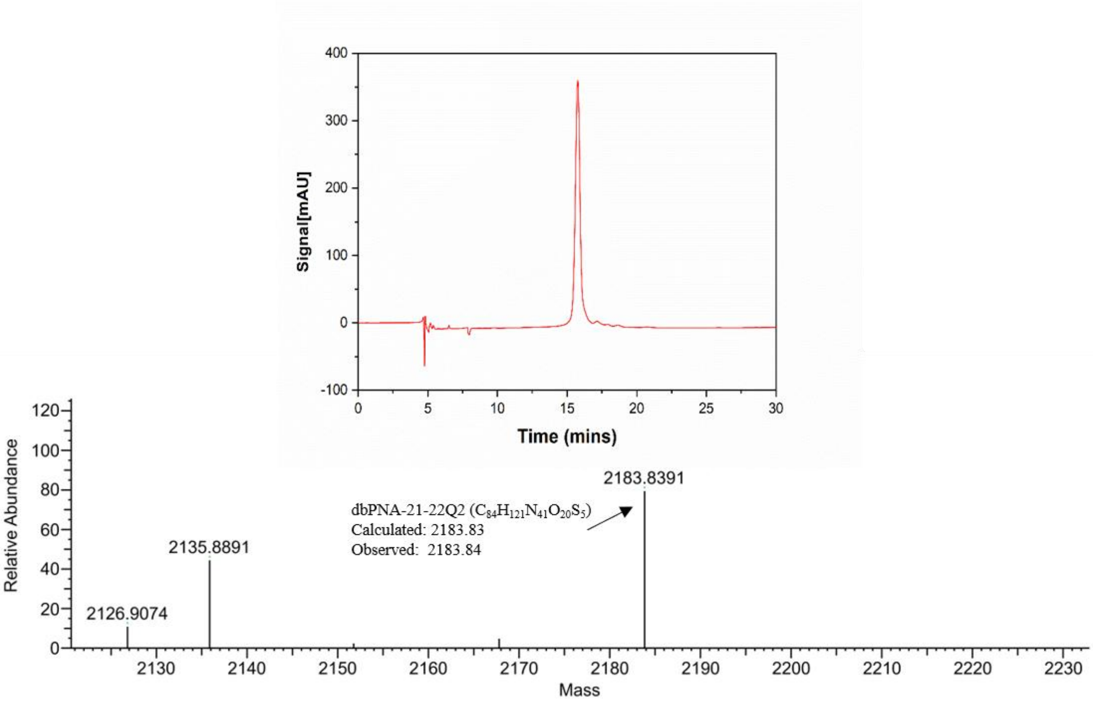


**Figure S6. HPLC and LC/MS data of daPNA21-22Q2.**

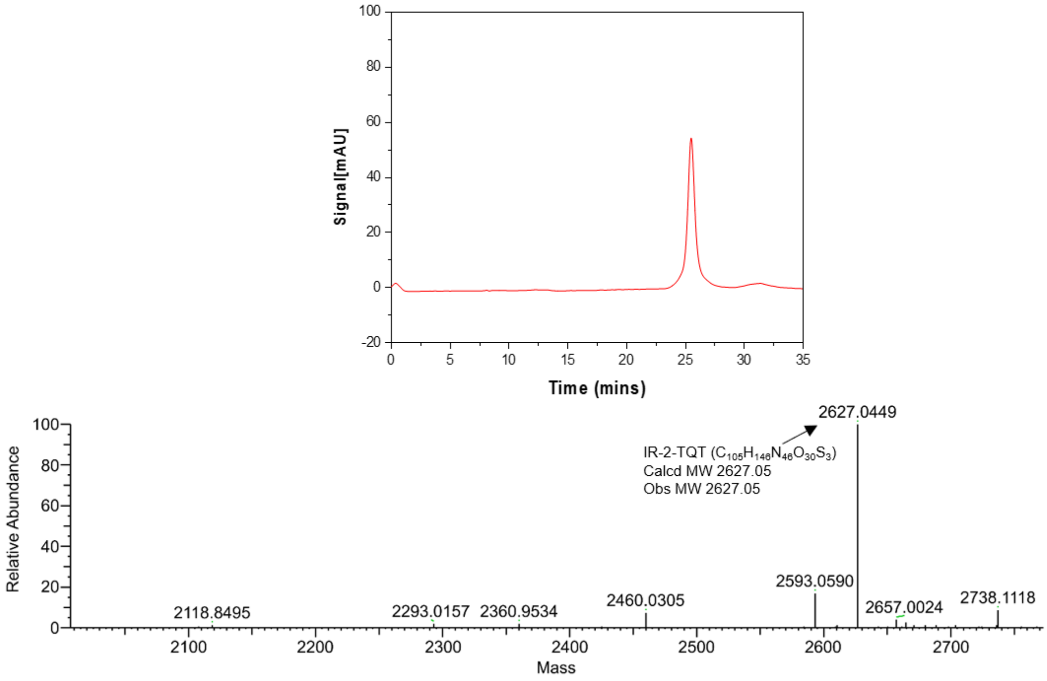


**Figure S7. HPLC and LC/MS data of IR-2-TQT.**

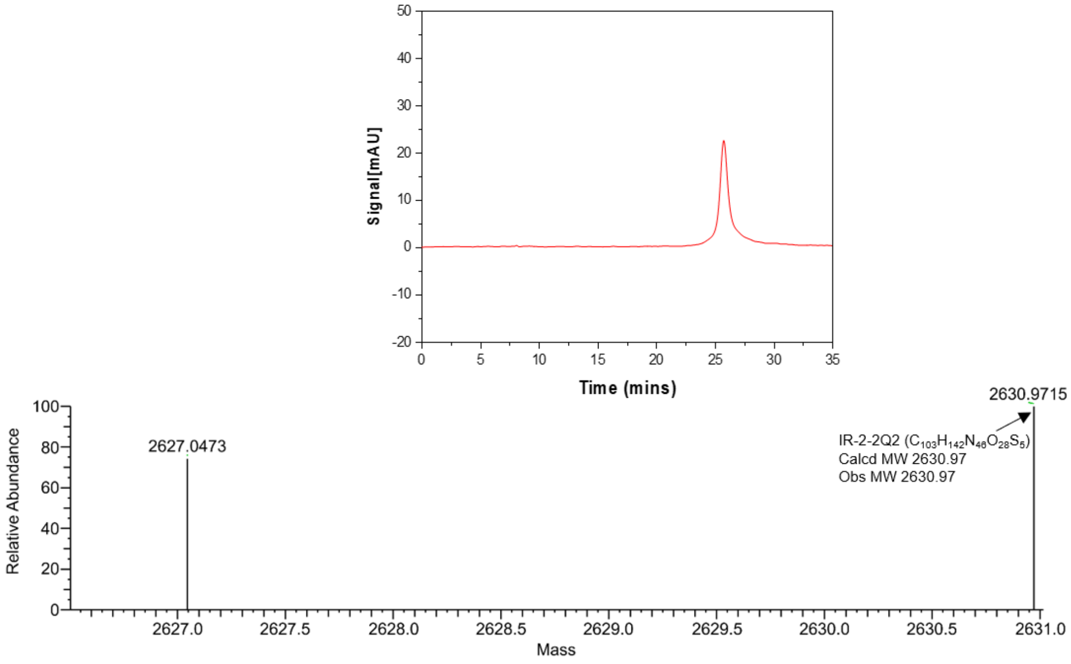


**Figure S8. HPLC and LC/MS data of IR-2-2Q2.**

### Table S2. DNA and RNA oligonucleotides used in this study.

| Oligomer | Label | Sequences (5’-3’) | DNA/RNA |
|---|---|---|---|
| rHP2-iCy3 | Cy3 | UAGAGAGAGAAAGUU**[Cy3]**CGACUUUCUCUCUCUA | RNA |
| rHP2-biotin | biotin | **Biotin**-UAGAGAGAcAAAGUUUCGACUUUgUCUCUCUA | RNA |
| Pre-miR-21 | biotin | **Biotin**-UAGCUUAUCAGACUGAUGUUGACUGUUGAAUCUCA UGGCAACACCAGUCGAUGGGCUGU | RNA |
| PH-v-biotin | biotin | AGUAGAAACAAGGGUGUUCGCACCCUGCUUUUGCU-**Biotin** | RNA |
| PH-v-Cy5 | Cy5 | AGUAGAAACAAGGGUGUUCGCACCCUGCUUUUGCU-**Cy5** | RNA |
| pDL-631 | / | CACTCCCAGTTCAATTACAG | DNA |
| pDL-3431 | / | TCTTATCATGTCTGCTCGAA | DNA |

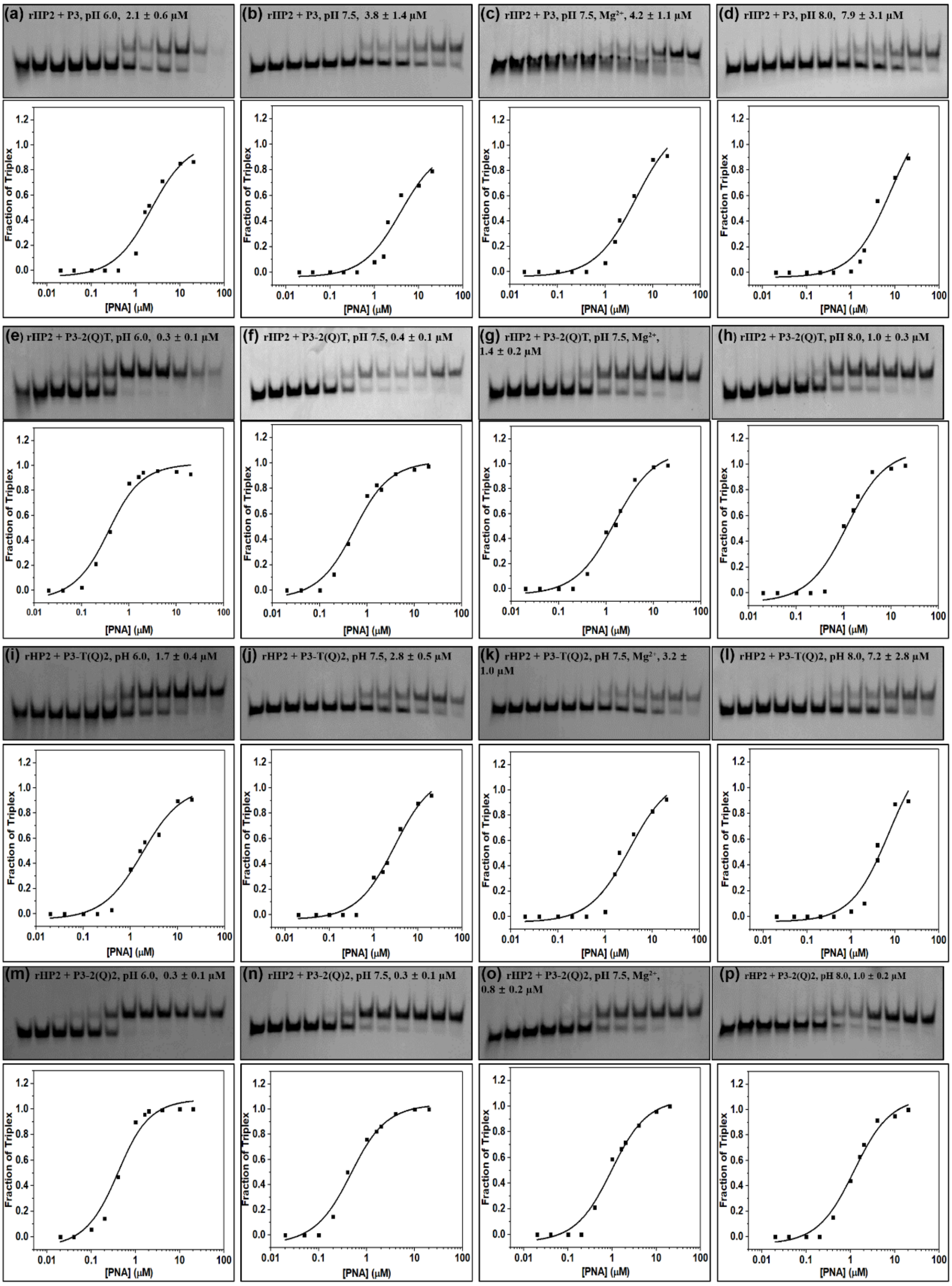


**Figure S9. PAGE characterization of PNAs designed for targeting rHP2.** Nondenaturing PAGE data for dbPNA P3, P3-2QT, P3-TQ2 and P3-2Q2. The incubation buffer is 200 mM NaCl, 0.5 mM EDTA and 20 mM HEPES. **(a-d)** The representative PAGE and fitting curves of P3 binding to rHP2 RNA construct at pH 6.0, pH 7.5, pH 7.5 with 2mM $Mg^{+}$, and pH 8.0, respectively. **(e-h)** The data of P3-2QT with rHP2 at various conditions. **(i-l)** The data of P3-TQ2 with rHP2 at various conditions. **(m-p)** The data of P3-2Q2 with rHP2 at various conditions. The PNA concentrations in lanes from left to right are 0, 0.02, 0.04, 0.1, 0.2, 0.4, 1, 1.6, 2, 5, 10, and 20 μM, respectively.

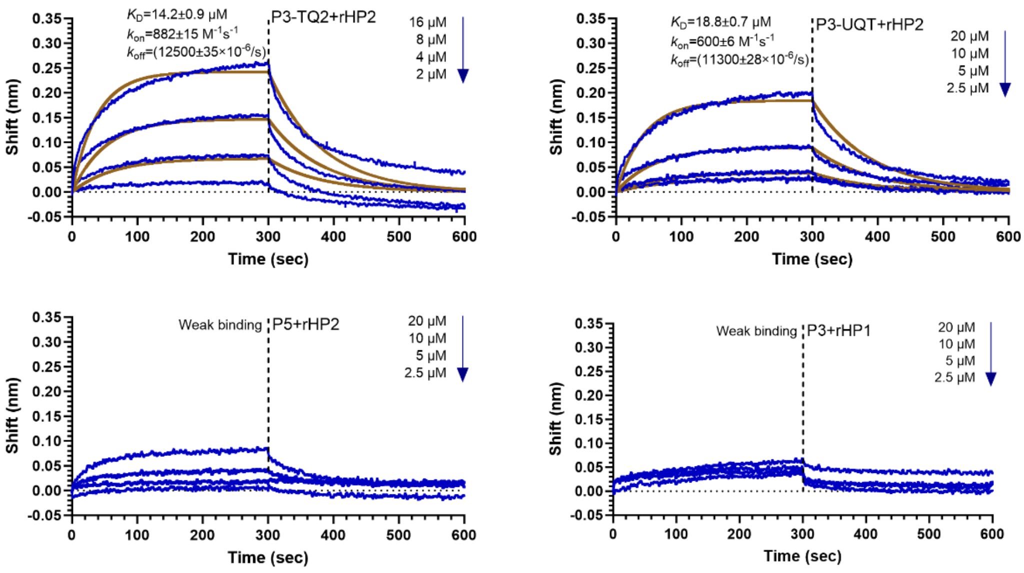


**Figure S10.** Bio-layer interferometry assay data for dbPNA binding to rHP2 and rHP1. The experimental data and fitting curves are shown in blue and brown, respectively. The final concentrations of the dbPNA are 20, 10, 5, and 2.5 μM from top to bottom except for P3-TQ2. The concentrations of P3-TQ2 are 16, 8, 4, and 2 μM.

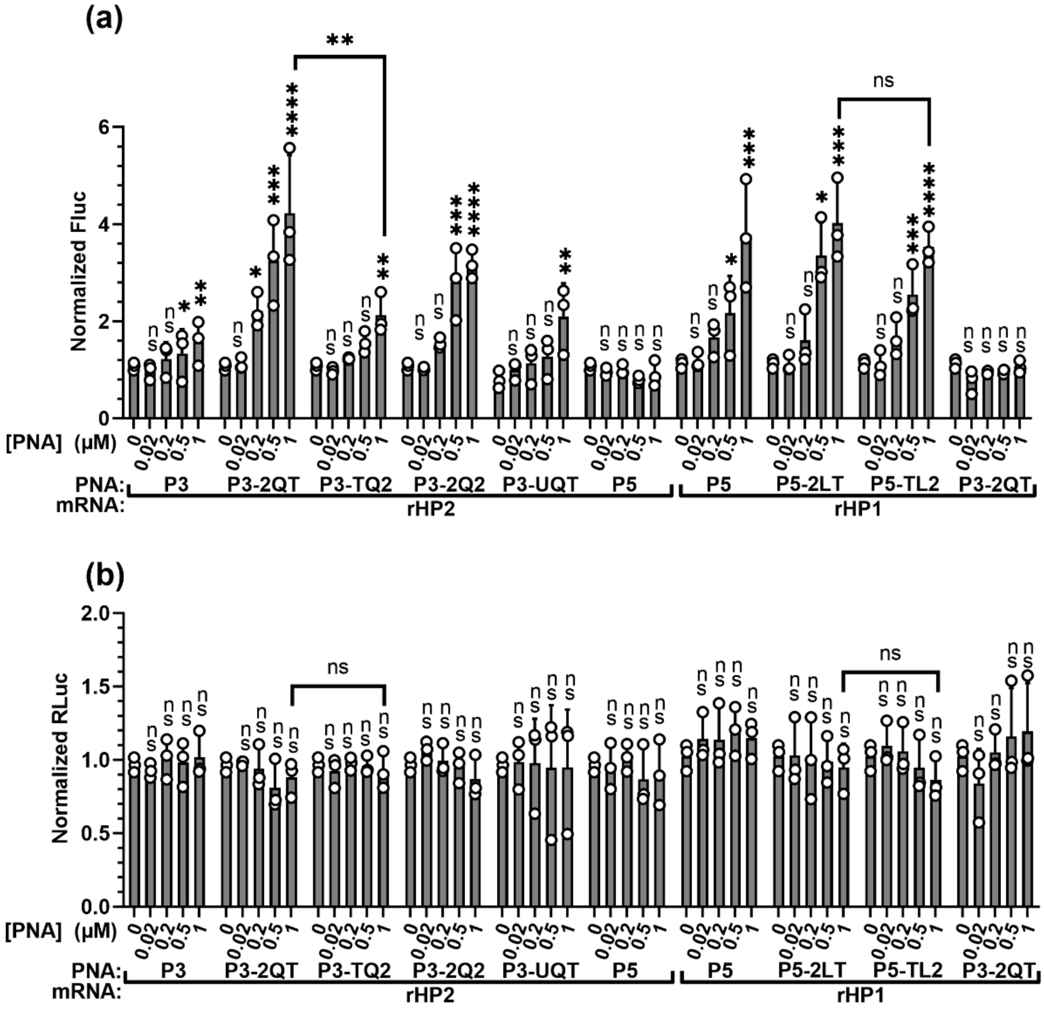


**Figure S11: Normalized cell-free dual-luciferase assay data for mRNAs rHP1 and rHP2.** Normalized Fluc (a) and Rluc (b) activities in cell-free dual-luciferase reporter assay were shown. The data were analyzed by GraphPad Prism 9.3 and calculated by an ordinary one-way analysis of variance (ANOVA) using Dunnett's multiple comparisons test against the mean of mRNA or plasmid alone group. The error bars represent ± S.D. * $P < 0.05$, ** $P < 0.01$, *** $P < 0.001$, **** $P < 0.0001$, ns: not significant.

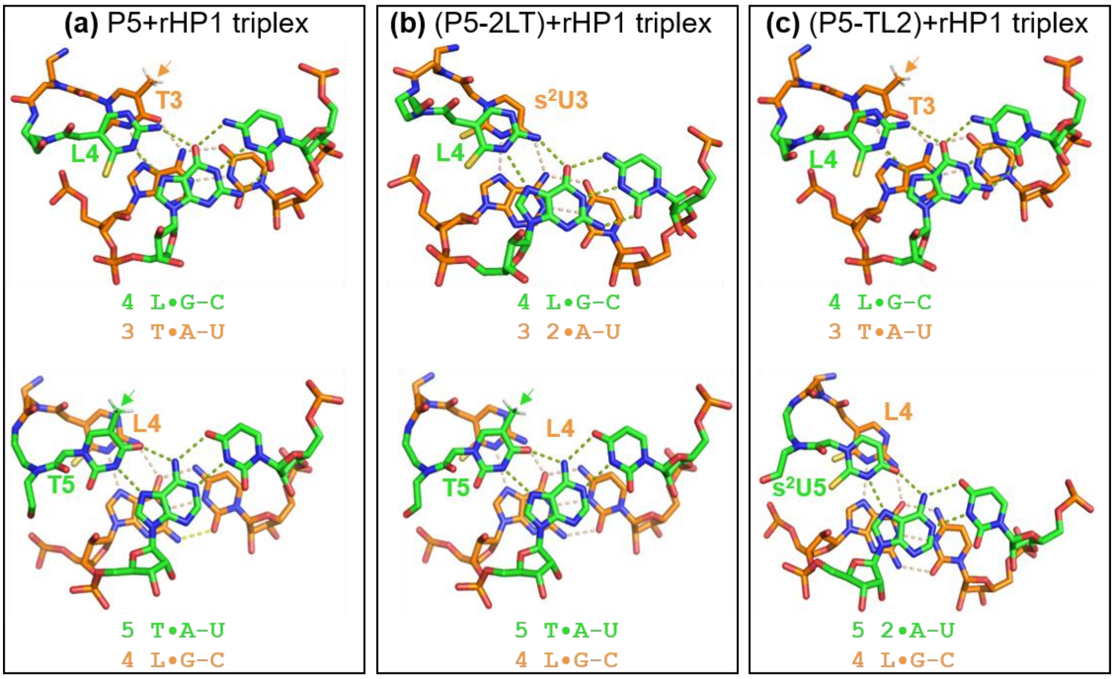


**Figure S12: Potential stacking patterns of PNA·RNA$_2$ base triples of dbPNAs targeting rHP1 involving PNA segments TLT/2LT/TL2.** The PNA strand is shown on the top left of each panel. (a) Stacking patterns involving residues 3 to 5 of P5 ($NH_2$-TLTLTTTL-$CONH_2$) targeting rHP1. (b) Stacking patterns involving residues 3 to 5 of P5-2LT ($NH_2$-TL2LTTTL-$CONH_2$) targeting rHP1. (c) Stacking patterns involving residues 3 to 5 of P5-TL2 ($NH_2$-TLTL2TTL-$CONH_2$) targeting rHP1. The hydrogen atoms of the methyl groups of T and Q are shown and indicated by an arrow. The stacking patterns were visualized by Pymol under orient (Vis) setting.

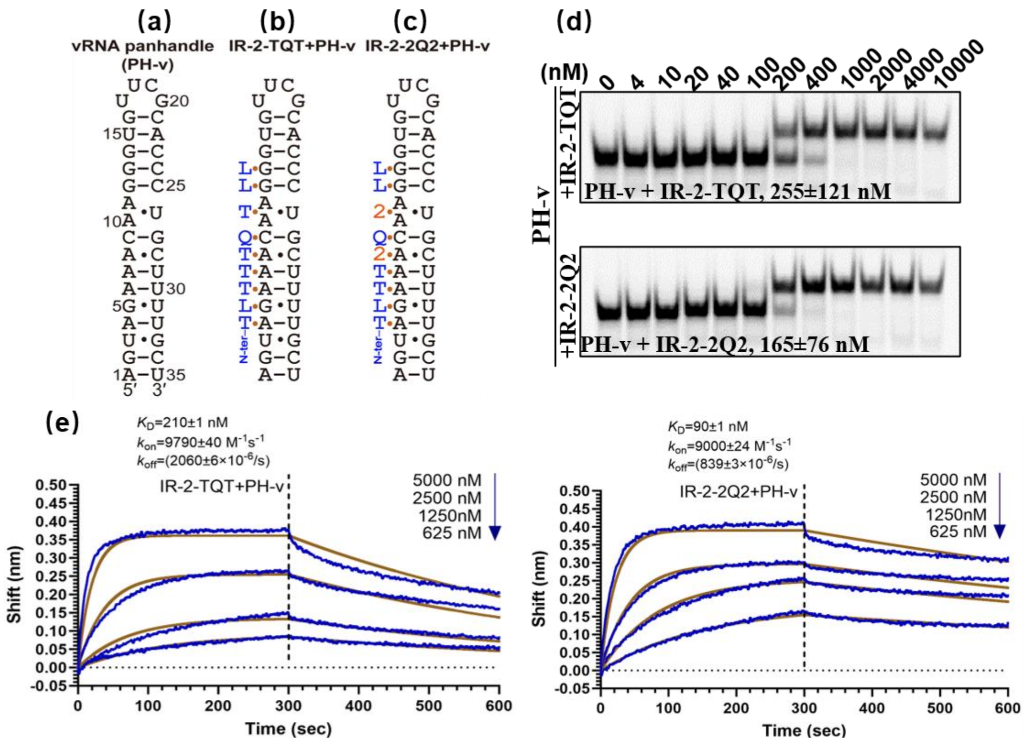


**(f)** **The binding parameters determined by bio-layer interferometry and PAGE.**

| PNA | RNA | PAGE | bio-layer interferometry | | | |
|---|---|---|---|---|---|---|
| | | $K_D$ (nM) | $K_D$ (nM) | $k_{on}$ ($M^{-1}$ $s^{-1}$) | $k_{off}$ (×$10^{-6}$ $s^{-1}$) | $R^2$ |
| IR-2-TQT | PH-v | 255±120 | 210±1 | 9790±41 | 2060±6 | 0.98 |
| IR-2-2Q2 | PH-v | 165±76 | 90±1 | 9000±24 | 839±3 | 0.99 |

**Figure S13. Schematics and binding properties for dbPNAs binding to the panhandle structure of influenza A virus (PH-v). (a-c)** Schematic of PH-v structure and triplexes formed with dbPNAs IR-2 and IR-2-2Q2. **(d)** Representative non-denatured PAGE data for IR-2-TQT and IR-2-2Q2 binding to 3′ Cy3-labeled PH-v RNA construct. **(e)** Representative BLI data for IR-2-TQT and IR-2-2Q2 binding to 3′ biotin-labeled PH-v RNA construct with a final concentration: 0, 625, 1250, 2500 and 5000 nM. **(f)** Binding parameters determined by non-denatured PAGE and BLI. The incubation buffer for each assay was 200 mM NaCl, 0.5 mM EDTA, 20 mM HEPS, pH=7.5.

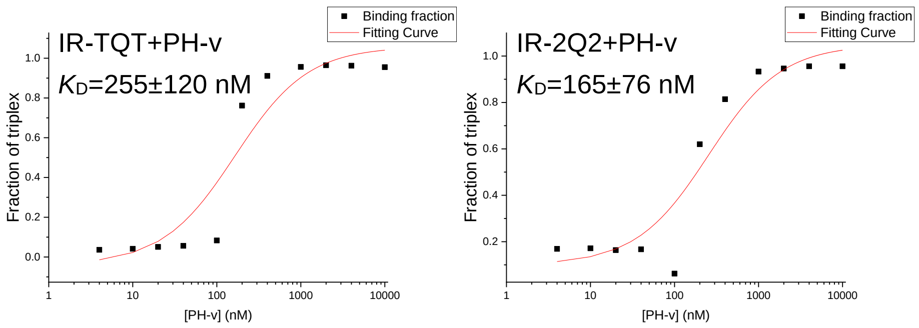


**Figure S14:** Fitting curve of PAGE results for dbPNAs binding to PH-v.

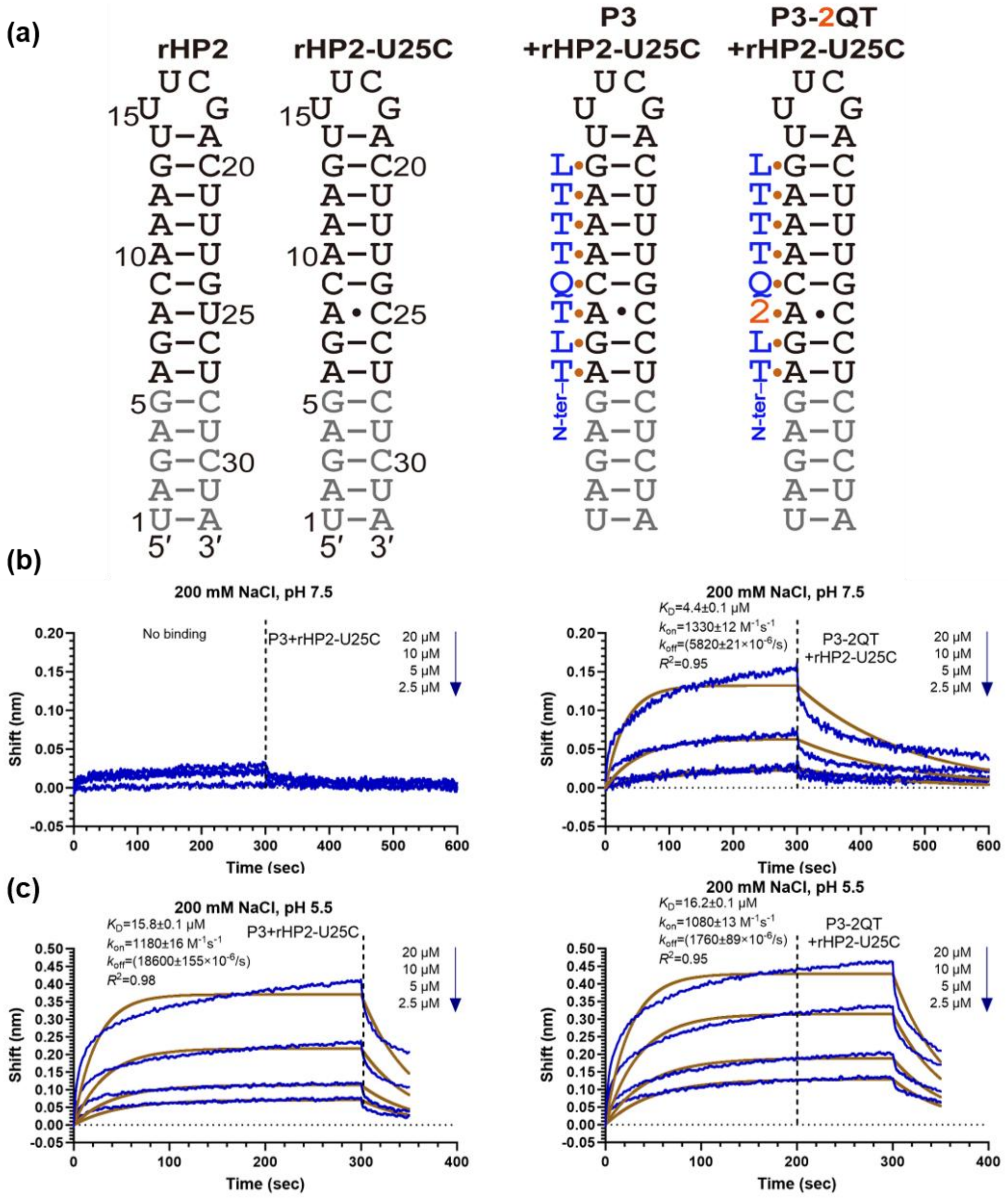


**Figure S15: Schematics and BLI binding data for dbPNAs binding to rHP2-U25C.** The experimental data and fitting curves are shown in blue and brown, respectively. The final concentrations of the dbPNA are 20, 10, 5 and 2.5 μM from top to bottom.

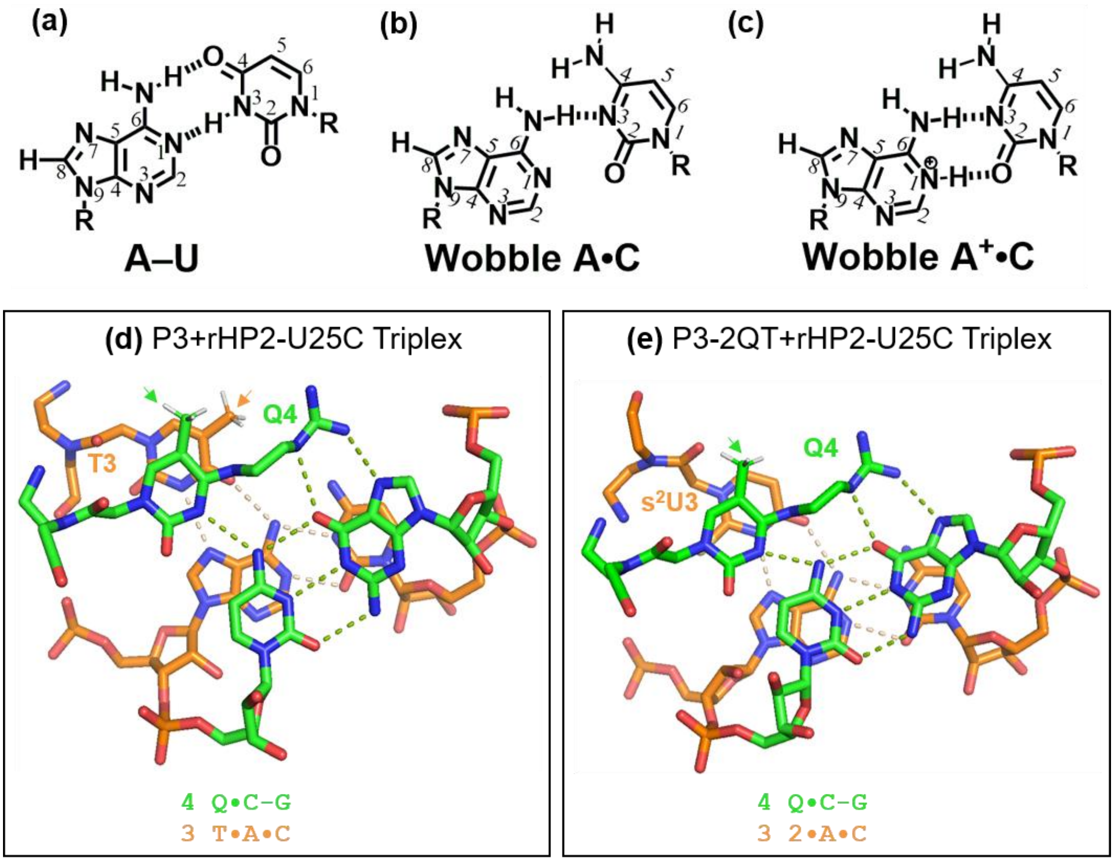


**Figure S16: Chemical structure comparison of A-U and A·C pairs and potential stacking patterns of PNA·RNA₂ base triples of dbPNAs targeting rHP2-U25C.** The PNA strand is shown on the top left of each panel. (a) Stacking patterns involving residues 3 to 4 of P3 ($NH_2$-TLTQTTTL-$CONH_2$) targeting rHP2-U25C. (b) Stacking patterns involving residues 3 to 4 of P3-2QT ($NH_2$-TL2QTTTL-$CONH_2$) targeting rHP2-U25C. The hydrogen atoms of the methyl groups of T and Q are shown and indicated by an arrow. The stacking patterns were visualized by Pymol under orient (Vis) setting.
